# The bacterial toxin ExoU requires a host trafficking chaperone for transportation and to induce necrosis

**DOI:** 10.1101/2020.11.04.367706

**Authors:** Vincent Deruelle, Stéphanie Bouillot, Viviana Job, Emmanuel Taillebourg, Marie-Odile Fauvarque, Ina Attrée, Philippe Huber

## Abstract

*Pseudomonas aeruginosa* is a causative agent in nosocomial infections, notably in ventilated or cystic fibrosis patients. *P. aeruginosa* isolates expressing the phospholipase ExoU, an effector of the type III secretion system, are the most pathogenic in clinics. Here, using a genome-wide screen, we discovered that ExoU toxicity requires DNAJC5, a host chaperone, to exert its necrotic activity. DNAJC5 is involved in an unconventional secretory pathway for misfolded proteins involving anterograde vesicular trafficking. DNAJC5-deficient human cells or *Drosophila* flies knocked-down for the DNAJC5 orthologue were largely resistant to ExoU virulence. ExoU colocalized with DNAJC5-positive vesicles in the host cytoplasm. DNAJC5 mutations preventing vesicle trafficking - identified in adult neuronal ceroid lipofuscinosis, a human congenital disease - inhibited ExoU-dependent cell lysis. These results suggest that, once injected into the host cytoplasm, ExoU docks to DNAJC5-positive secretory vesicles to reach the plasma membrane, where its phospholipase activity is triggered by binding to PI(4,5)P2.

## Introduction

In most instances, bacterial toxins require one or more host factors to exert their toxicity. These factors can be receptors, binding partners inducing structural modifications or even entire host cellular pathways that are hijacked for bacterial toxicity purposes. The requirement for host-cell mechanisms protects bacteria from self-toxicity and takes advantage of efficient molecular mechanisms developed by eukaryotic cells to alter cellular functions. This rule applies to the toxins secreted by *Pseudomonas aeruginosa*, a Gram-negative opportunistic pathogen.

*P. aeruginosa* is a leading cause of severe nosocomial infections. It is a causative agent of pneumonia, urinary tract infections, bacteraemia, abscesses, as well as burn and eye infections. *P. aeruginosa* infections are frequent in ventilated and cystic fibrosis patients, and have a particularly high fatality rate following infection in these conditions^1–3^. The high mortality rate recorded is also due to acquired resistance to antibiotics, which is a major issue in the clinical management of *P. aeruginosa* infections^4–6^.

*P. aeruginosa* uses a multi-target strategy to infect host cells, employing a combination of virulence factors. One of these factors is the type 3 secretion system (T3SS), the effectors of which are known to be the most potent toxins in acute *P. aeruginosa* infections^2,7^. The T3SS consists of a syringe-like apparatus which injects toxins into the cytosol of host cells. Four effectors have been identified: ExoU, ExoS, ExoT and ExoY, having their cognate co-activation host factors: phosphatidylinositol-4,5-bisphosphate [PI(4,5)P2] and ubiquitin for ExoU, 14-3-3 adaptor protein for ExoS and ExoT, and filamentous actin for ExoY^8–14^.

ExoU and ExoS are mutually exclusively expressed in bacterial strains ^2^. ExoU-positive bacteria represent 28-48% of *P. aeruginosa* clinical isolates, and are found in the most severe pathological cases and produce the most dramatic lesions^2,7,15^. Furthermore, ExoU-positive strains have been associated with increased multidrug resistance in several clinical studies^16–19^.

ExoU is a phospholipase A2 (PLA2) inducing plasma membrane rupture and rapid cell necrosis ^20,21^. Its activity is enhanced by binding to ubiquitin and to the lipid PI(4,5)P2, a lipid present in the inner leaflet of the plasma membrane^10–14^. However, several aspects of ExoU activation and trafficking in host cells remain elusive. Here, we searched for other host factors required for full ExoU toxicity using a genome-wide screening approach and we identified DNAJC5 as a necessary cofactor for its trafficking in host cells.

## Results

### ExoU requires DNAJC5 for host cell lysis

To identify host genes involved in ExoU cytotoxicity, we performed a genetic screen using CRISPR-cas9 technology. A549 pneumocytic cells were transduced with a lentiviral library of guide-RNAs (gRNAs), targeting 18,053 genes (four gRNAs per gene). The cells were subjected to three rounds of infection with the *P. aeruginosa* strain PA14, known to induce cell necrosis via ExoU secretion (Fig. 1a). Each infection round was stopped by adding antibiotics after 90 min of infection, and each round of infection resulted in approximately 70% of cell death. This experimental design aimed at selecting resistant cells to ExoU-induced necrosis putatively carrying a mutated human gene required for ExoU necrotizing activity. The gRNA sequences in surviving A549 cells were identified by next-generation sequencing and the number of reads for each gRNA was compared to the number in the library of uninfected library. Three independent replicates were performed and a statistical analysis revealed a significant enrichment for gRNAs targeting only one gene: the gene encoding DNAJC5 (also known as cysteine string protein α; CSPα)(Fig. 1b).

**Figure 1:**
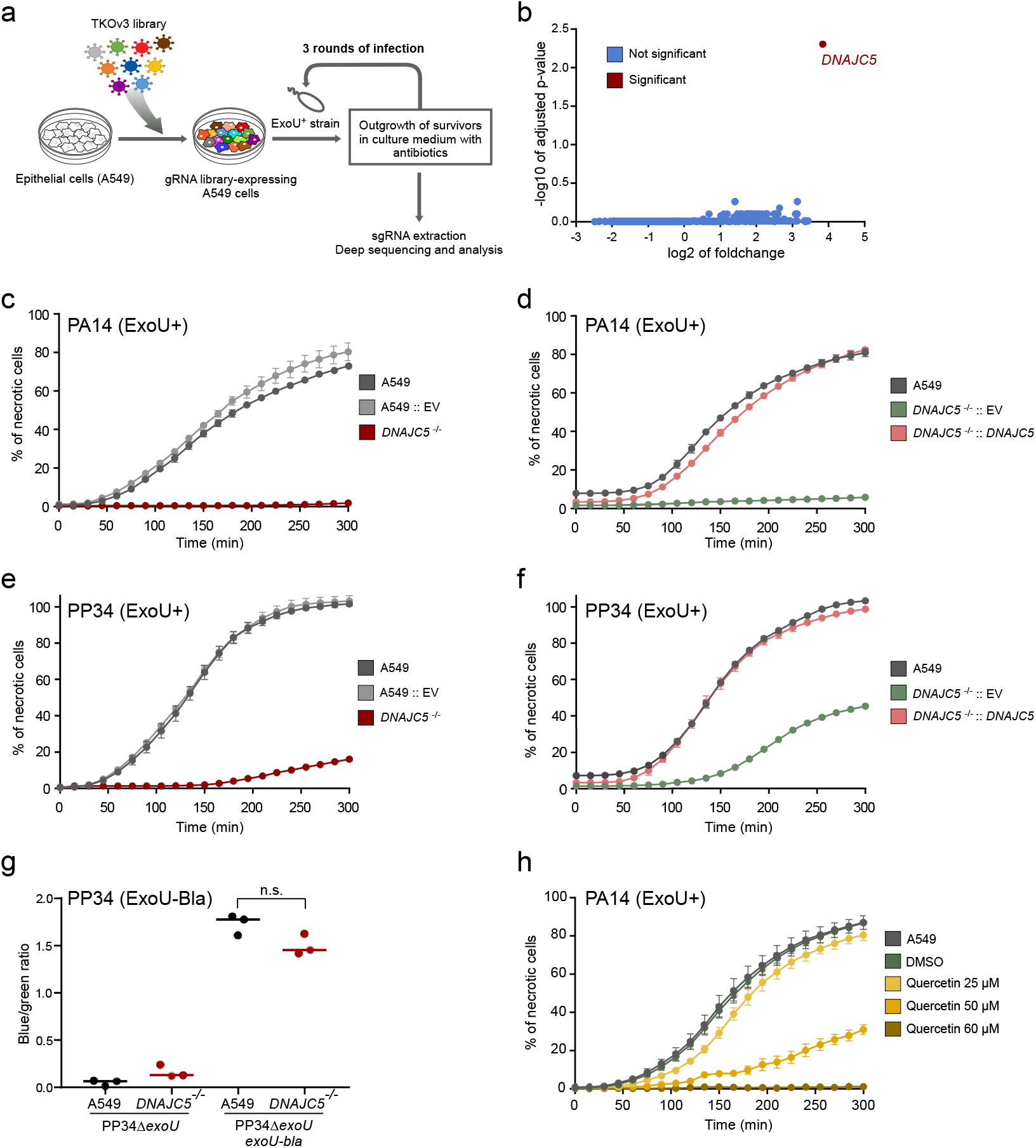
DNAJC5 is required for ExoU cytotoxicity. **a**. Screening process to identify host genes required for ExoU toxicity. A gRNA library (TKOv3, four gRNAs per gene) was constructed for A549 human epithelial cells. Cells were subjected to three 90-min rounds of infec-tion with the ExoU+ PA14 strain for in triplicates. Infection was stopped by washing and adding antibiotics. gRNAs were then amplified by PCR and submitted to deep sequencing. **b**. Analysis of sequencing data. Statistical analysis of gRNA amplification in infected vs uninfected conditions. gRNAs targeting the DNAJC5 gene were the only ones significantly enriched in the screen. **c**. Cytotoxicity assay. A clonal population of DNAJC5^−/−^ A549 cells, native A549 cells or cells transfected with an empty vector (EV) were infected in 6-replicates with PA14 and necrosis was monitored by propidium iodide incorporation, recorded by time-lapse microscopy. Results are represented as the mean percentage (+/−SD) of necrotic cells. **d**. Cytotoxicity assay with A549 native cells. A549 DNAJC5^−/−^ cells complemented with DNAJC5 (DNAJC5^−/−^::DNAJC5), or the mock-complemented control, were infected with PA14. **e, f**. Cytotoxicity assays similar to c,d, but cells were infected with the ExoU+ PP34 strain. **g**. T3SS-dependent injection of ExoU in A549 cells. A549 cells or DNAJC5^−/−^ A549 cells were infected with PP34Δ*exoU* bacteria complemented with exoU fused to the ß-lactamase gene (*bla*) or infected with uncomplemented PP34Δ*exoU*. Cells were loaded with CCF2, a Bla fluorescent substrate which shifts from green to blue fluorescence upon processing by the enzyme. Fluorescence was measured at 4 hpi on both channels and results are expressed as a blue/green ratio. **h**. Cytotoxicity assay. Increasing concentrations of quercetin were added to A549 cells in the presence of PA14 (ExoU+) at an MOI of 20. Necrosis was monitored by PI incorporation and recorded by time-lapse microscopy.

DNAJC5 is a ubiquitous cytoplasmic protein located at the surface of late endosomes (LEs). It functions as a co-chaperone in association with Hsc70 or Hsp70, which play a central role in protein homeostasis^22–24^. DNAJC5 is also required for an unconventional protein secretion pathway^25,26^, recently described as Misfolded-Associated Protein Secretion (MAPS)^27^. In this process, misfolded cytosolic proteins are translocated into DNAJC5+ LEs near the endoplasmic reticulum, which are then transported to the plasma membrane^28,29^.

Eventually, fusion of the vesicles with the plasma membrane allows the elimination of misfolded proteins directly into the extracellular milieu^27^. Alternatively, the vesicles can produce exosomes containing the misfolded proteins in the extracellular milieu^30^. This secretion process has been widely described in neurons, where alteration of MAPS in neurons has been linked to several neurological disorders^26,31–35^.

To confirm the role of DNAJC5 in ExoU-dependent cytotoxicity, we generated independent DNAJC5^−/−^ A549 cells using CRISPR-cas9 technology and one of the gRNA targeting *DNAJC5* in the library (Supplementary Fig. 1a). A clonal population was selected, lacking four bases from the coding sequence of both *DNAJC5* alleles. The DNAJC5^−/−^ cells were subjected to a cytotoxic test, after infection with PA14 in the presence of propidium iodide (PI) to detect necrotic cells. PI incorporation was monitored by automated time-lapse microscopy (Fig. 1c). The proportion of native A549 cells exhibiting a necrotic phenotype increased with time and reached > 75% at 5 h post-infection (pi); in contrast, DNAJC5^−/−^ cells exhibited no PI incorporation. PA14 lytic capacity was restored when *DNAJC5* expression was rescued in DNAJC5^−/−^ cells (DNAJC5^−/−^::DNAJC5)(Fig. 1d). Similar experiments were performed with the clinical strain PP34, isolated from bacteraemia and secreting high amounts of ExoU. In these experiments, some necrosis (18%) was observed at late time points in DNAJC5^−/−^ cells, while 100% of A549 cells were necrotic (Fig. 1e). As with PA14, PP34 infection of DNAJC5^−/−^::DNAJC5 cells restored a full ExoU cytotoxicity (Fig. 1f).

To determine whether DNAJC5 contributes directly to toxin activity, or is required for T3SS-dependent injection, we infected cells with bacteria secreting ExoU fused to β-lactamase (ExoU-Bla). This reporter system was previously used to monitor ExoU delivery into cells^36^. Host cells were pre-loaded with a fluorescent substrate of Bla (CCF2) used for fluorescence resonance energy transfer (FRET) experiments. Uncleaved CCF2 produces a green fluorescence, whereas the cleaved CCF2 emits a blue fluorescence. Infection of A549 or DNAJC5^−/−^ cells with ExoU-Bla-secreting bacteria produced similar ratios of blue/green fluorescence (Fig. 1g), indicating that the absence of DNAJC5 did not alter T3SS injection *per se*.

As a complementary demonstration that DNAJC5 is required for ExoU toxicity, we used a DNAJC5 inhibitor. Quercetin is a natural product inducing DNAJC5 dimerization at high concentrations, leading to its inactivation^37^. Upon application of quercetin to cell cultures, a dose-dependent inhibition of ExoU-induced A549 cell lysis was observed (Fig. 1h), confirming the essential role of DNAJC5 in ExoU intoxication.

We next examined whether DNAJC5 also contributes to the activity of other toxins delivered by *P. aeruginosa*. To do so, we first performed a cytotoxicity assay using bacteria (CHAΔ*exoT*) secreting ExoS (and not ExoU or ExoT) through the T3SS. The toxic activity of ExoS results in dismantling of the actin cytoskeleton, hence provoking cell rounding ^2^. The similar kinetic profiles recorded for the two cell lines (Supplementary Fig. 2a,b), indicate that the action of ExoS in host cells does not require DNAJC5. This result also further confirmed that T3SS injection is unaffected in DNAJC5^−/−^ cells.

We subsequently assayed how DNAJC5 deficiency affected toxicity of the pore-forming toxin ExlA, secreted by *P. aeruginosa* strains lacking T3SS^38^. Like ExoU, ExlA is a necrotizing toxin. Identical intoxication curves were recorded for A549 and DNAJC5^−/−^ cells (Supplementary Fig. 2c), indicating that DNAJC5 is not involved in ExlA-dependent cell lysis. Likewise, DNAJC5 seems to be specifically required for ExoU necrotizing activity in host cells.

Taken together, these results demonstrate that DNAJC5 is specifically required for ExoU toxicity in host cells.

### Lack of DNAJC5 decreases the virulence of ExoU-positive *P. aeruginosa*’s strains in vivo

DNAJC5 is an evolutionary conserved protein. Animals in which the *DNAJC5* orthologue gene was inactivated (mice, *Caenorhabditis elegans* and *Drosophila melanogaster*) all rapidly died after birth from neurological disorders^39–41 39–41^. These models are consequently unsuitable for use in infection assays. To overcome this lack of in vivo model, we generated *Drosophila* in which the *DNAJC5* orthologue *Cystein string protein* (*Csp*) was conditionally knocked-down (KD) by expressing a silencing long double-strand RNA (dsRNA) or a short hairpin RNA (shRNA) in two independent fly lines (generation of the *Csp*-KD flies is shown in Supplementary Fig. 3a,b). Flies were infected by pricking the thorax with a thin needle previously dipped into a bacterial suspension^42^ (Fig. 2a). This infection model can be used to measure the impact of ExoU on fly death, as shown by the difference in survival curves following infection with a wild-type strain of *P. aeruginosa* expressing high levels of ExoU (PP34) and its isogenic mutant (PP34Δ*exoU*) (Supplementary Fig. 3c). Survival curves for control and *Csp*-KD *Drosophila* infected with PP34 were strikingly different (Fig. 2b,c), whereas the survival curves for *Csp*-KD and *Drosophila* mock-infected with PBS were not statistically different. These data indicate that DNAJC5/CSP is required for full ExoU-dependent *P. aeruginosa* virulence in vivo.

**Figure 2:**
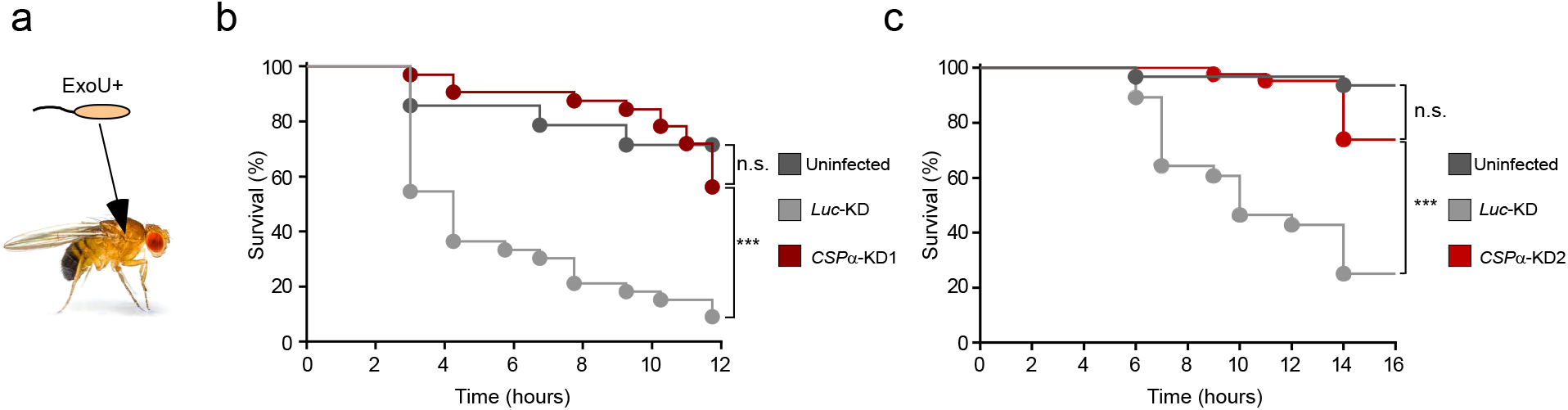
ExoU toxicity in Drosophila requires CSP (DNAJC5 orthologue). **a**. Drosophila were infected with PP34 (ExoU+) by pricking the thorax with a thin dipped in bacterial suspension. **b, c**. Flies expressing RNAi transgenes targeting either the firefly *luciferase* gene (*Luc*-KD) or the *Csp* gene (*Csp*-KD1 and *Csp*-KD2). Mock-infected *Csp*-KD1 (n = 40) or *Csp*-KD2 (n = 51) flies with PBS were included as negative control (uninfected; n = 30 and 47, respectively). Fly survival was recorded over 12 hours and data are represented as Kaplan-Meyer curves. Statistical differences were established using the Log-Rank test. n.s., not significant; ***, p < 0.001.

### ExoU partly localizes in DNAJC5-positive vesicles

To determine the localization of ExoU in infected cells, we used a *P. aeruginosa* strain (CHAΔ*exoSexoT::exoU*^*S142A*^, hereafter CHA-*exoU*^*S142A*^) that secretes a catalytically-inactive non-lytic mutant, ExoU^S142A 43^. Cells were infected with this strain and soluble and membrane fractions were prepared from lysates of A549 infected cells. ExoU was mainly detected in the membrane fractions, with only trace amounts present in soluble fractions (Fig. 3a). This distribution indicates that following injection, ExoU binds to membranes rather than remaining free in the cytosol. Equivalent amounts of ExoU were detected in DNAJC5^−/−^ and DNAJC5^−/−^::DNAJC5 membrane fractions, demonstrating that DNAJC5 is not required for ExoU docking to membranes. To gain further insight into the subcellular localization of ExoU, we performed immunofluorescence experiments and observed cells by confocal microscopy. In infected A549 and DNAJC5^−/−^ cells, ExoU displayed similar particulate labelling in the cytoplasm of both cell types (Fig. 3b). Thus, once ExoU is injected into cells, it binds to specific cytoplasmic structures in a DNAJC5-independent manner.

**Figure 3:**
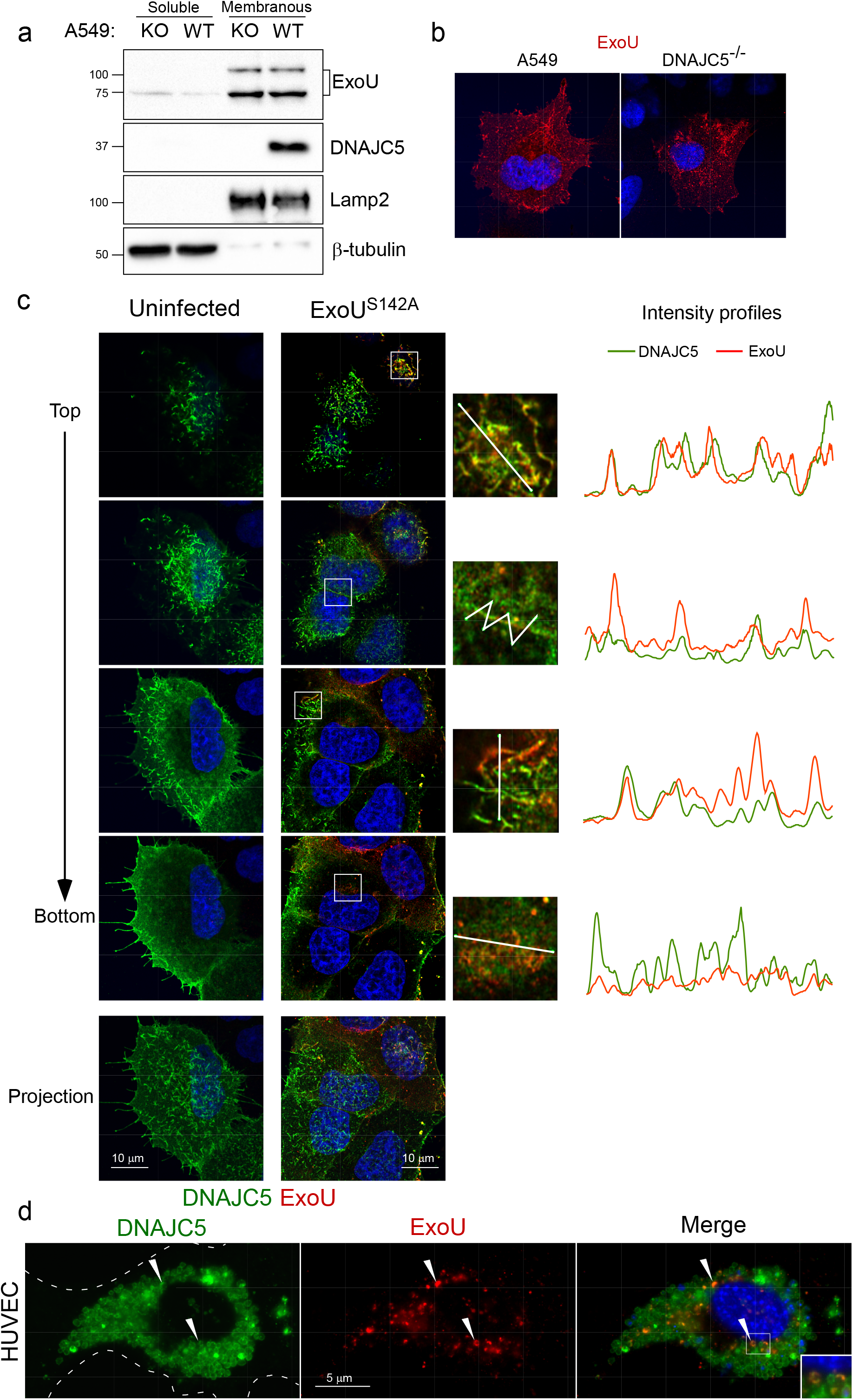
ExoU partly localizes in DNAJC5-positive vesicles. **a.** Fractionation of DNAJC5 ^−/−^ (KO) and DNAJC5^−/−^::DNAJC5 (WT) cells infected with a *P. aeruginosa*’s strain (CHA-*exoU*^*S142A*^) secreting a catalytically inactive ExoU mutant through its T3SS.Cells were harvested at 4 hpi and their soluble and membrane fractions were prepared. Western blots were performed on fractions using anti-ExoU, DNAJC5, Lamp2 (a late endosome - lysosome marker) and ß-tubulin (a cytosolic marker) antibodies. Thehigher molecular weight bands revealed by the ExoU antibody represent the ubiquitinylated form of ExoU^66^. **b**. ExoU immunofluorescence staining of A549 and DNAJC5^−/−^ cells infected with CHA-*exoU*^*S142A*^. A single representative z-section is shown. **c left.** DNAJC5-FLAG immunofluorescence signals (green) in uninfected DNAJC5^−/−^::DNAJC5 cells. Four z-sections obtained by confocal microscopy are shown from top to bottom. A z-projection is also shown below. Nuclei were counterstained in blue. **c right**. DNAJC5-FLAG (green) and ExoU (red) immunofluorescence signals in DNAJC5^−/−^::*DNAJC5* cells infected with CHA-*exoU*^*S142A*^. As for uninfected cells, four z-sections and a z-projection are shown. For each section, a region was enlarged and a bar was drawn (represented on the right) to establish an intensity profile for both green and red fluorescences, as shown. **d**. DNAJC5-GFP and ExoU localizations in transfected HUVEC infected with CHA-*exoU*^*S142A*^ on a wide-field microscopy image. Arrowheads show colocalization of both markers. The insert is an enlargement of themerged image, showing DNAJC5 and ExoU localization at the vesicle’s membrane.

To determine whether ExoU colocalizes with DNAJC5 in these intracellular structures, ExoU and DNAJC5-Flag were labelled in DNAJC5^−/−^::DNAJC5 cells infected with CHA-*exoU*^*S142A*^. Most DNAJC5-FLAG labelling was observed at the cellular periphery, associated with elongated vesicles, and at cell-cell junctions (Fig. 3c). Round DNAJC5+ vesicles were also present in the perinuclear region. These perinuclear DNAJC5+ LEs were also positive for the lysosome/late endosome marker Lamp2 (Supplementary Fig. 4), as previously reported for Cos7 cells^28^.

ExoU was associated with DNAJC5 in both elongated and round vesicles, as well as at cell-cell junctions, in the top and middle parts of the cell. However, in the lower part of the cell, the two labels were dissociated (Fig. 3c). Interestingly, the presence of ExoU had no impact on the subcellular localization of DNAJC5+ vesicles.

To allow a more detailed examination of DNAJC5/ExoU colocalization, we performed a similar experiment with endothelial cells (HUVECs), which have larger, mostly perinuclear, DNAJC5+ LEs. In these cells, both DNAJC5-GFP and ExoU were localized at the vesicle’s limiting membrane and were not intraluminal (Fig. 3d), confirming that ExoU is membrane-associated.

### Hsp70 and Hsc70 chaperones are dispensable for ExoU toxicity

As the T3SS does not accommodate folded proteins^44^, ExoU is probably delivered unfolded into host cells. Therefore, we reasoned that ExoU might need the chaperone activity of the DNAJC5-Hsc70/Hsp70 complex to recover its catalytic activity.

Three domains have been identified in DNAJC5 (Fig. 4a). A J domain, present in all DNAJ proteins, a cysteine string domain containing 14 cysteines and a C-terminal domain^32^. The J domain interacts with Hsc70/Hsp70 and enhances the ATPase activity of these chaperones^45^. Palmitoylation of cysteines in the central domain allows initial DNAJC5 anchoring at the surface of endosomes^46^. The C-terminal domain has been shown to associate with various proteins, including the vesicle-associated membrane protein (VAMP), a SNARE protein involved in the fusion of vesicles with the plasma membrane^47,48^.

**Figure 4:**
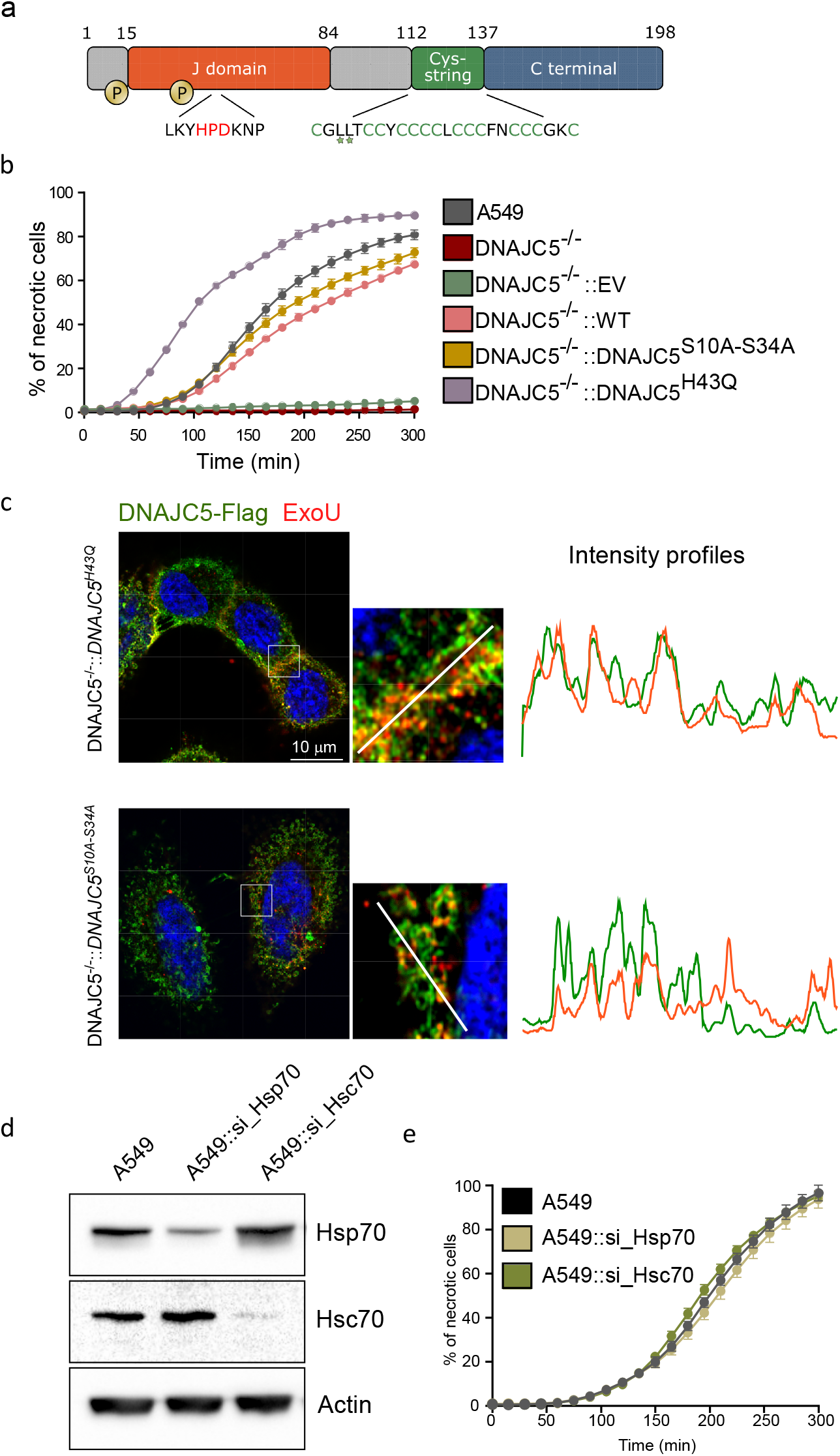
DNAJC5 co-chaperone activity is dispensable for ExoU toxicity. Domain organization of the human DNAJC5 protein. The following locations are highlighted: the two phosphory-lation sites (serine 10 and serine 34), the HPD motif for Hsc70/Hsp70 binding, and the cysteine-rich region con-taining the leucines L115 and L116, mutated in adult neuronal ceroid lipofuscinosis patients (stars). **b**. Effect of DNAJC5 mutations, H43Q and S10A-S34A, DNAJC5 mutations on ExoU toxicity. A549 cells, as well as DNAJC5^−/−^ cells complemented with either DNAJC5, DNAJC5H43Q, DNAJC5S10A-S34A or the empty vector (EV) were subject-ed to an infection assay with PA14. N = 6 fields per condition. **c**. DNAJC5-FLAG (green) and ExoU (red) immuno-fluorescence signals for DNAJC5^−/−^::DNAJC5H43Q and DNAJC5^−/−^::DNAJC5S10A-S34A cells infected with CHA-exoUS142A strain, which secretes a catalytically-inactive ExoU mutant. One z-section is shown. For each section, a region was enlarged and a bar was drawn (represented on the right) to establish the intensity profiles for both green and red fluorescence. **d, e**. Effect of decreased Hsc70 and Hsp70 expression on ExoU toxicity. A549 cells were transfected with siRNAs for Hsp70 (si_Hsp70) or Hsc70 (si_Hsc70) to knock down their expression. Knock-down was monitored by Western blot (**d**). KD cells were subjected to an infection assay (**e**), in the same conditions as in (**b**).

To determine whether the co-chaperoning role of DNAJC5 is linked to ExoU toxicity, we produced cells carrying several mutations to disrupt interactions between DNAJC5 and Hsp70/Hsc70, and infected them with ExoU+ *P. aeruginosa*. First, we complemented DNAJC5^−/−^ cells with DNAJC5^H43Q^, as this mutation was previously shown to disrupt DNAJC5-Hsc70/Hsp70 interaction^49^. DNAJC5^−/−^::DNAJC5^H43Q^ cells displayed higher sensitivity to *P. aeruginosa*-induced cell lysis than DNAJC5^−/−^::DNAJC5 (Fig. 4b), indicating that DNAJC5 interaction with Hsc70/Hsp70 is not required for ExoU activation, and instead delays ExoU-dependent cell lysis.

Shirafuji et al have reported that phosphorylation of two serines (S10 and S34) enhanced the co-chaperone activity of DNAJC5^50^. Furthermore, they showed that replacing the two serines by alanines (DNAJC5^S10A-S34A^) reduced DNAJC5 interaction with Hsp70. Therefore, we used DNAJC5^−/−^ cells complemented with DNAJC5 carrying both mutations in a cytotoxicity assay. ExoU displayed similar toxicity in DNAJC5^−/−^::DNAJC5 and DNAJC5^−/−^::DNAJC5^S10A-S34A^ cells(Fig. 4b), providing further evidence that DNAJC5’s co-chaperone activity is not required for ExoU activation. Interestingly, the H43Q and S10A-S34A mutations had no effect on subcellular localization of DNAJC5, nor on its colocalization with ExoU and Lamp2 (Fig. 4c and Supplementary. 4 and 5).

To definitively investigate the role played by Hsp70/Hsc70 in ExoU toxicity, we knocked-down each of these two proteins in A549 cells using siRNAs (Fig. 4d). Both KD cell lines were sensitive to PA14-induced lysis with a similar profile to native cells (Fig. 4e).

Based on these results, and despite the proven role of Hsc70/Hsp70 in protein exocytosis in MAPS^26^, these chaperones appear not to be required for ExoU intoxication.

### DNAJC5 escort of ExoU to the plasma membrane is required for toxicity

Having eliminated its co-chaperone role, we next investigated whether the trafficking activity of DNAJC5 was required for ExoU-dependent necrosis.

Adult neuronal ceroid lipofuscinosis is a neurodegenerative disease caused by mutations in *DNAJC5*^31,34,35^. Two mutations have been reported in patients: L115R and ΔL116 (stars in Fig. 4a). These mutations form protein oligomers^51,52^ that impair the trafficking of DNAJC5+ vesicles towards the plasma membrane, causing the accumulation of misfolded proteins in cells, and leading to progressive neuronal dysfunction^51^. When expressed in DNAJC5^−/−^ cells, DNAJC5^L115R^ and DNAJC5^ΔL116^localized in perinuclear vesicles, but not in vesicles at the cellular periphery (Supplementary Fig. 5), a feature previously reported in PC12 neuroblastic cells^51^. Furthermore, these DNAJC5+ vesicles were not labelled with Lamp2, which is associated with other perinuclear vesicles (Supplementary. 4). In both DNAJC5^−/−^::DNAJC5^L115R^ and DNAJC5^−/−^::DNAJC5^ΔL116^, ExoU colocalized with DNAJC5-FLAG (Fig. 5a,b). However, ExoU toxicity was severely diminished, recovering only minimally with DNAJC5^L115R^ or moderately with DNAJC5^ΔL116^ in these complemented cell lines (Fig. 5c).

**Figure 5.**
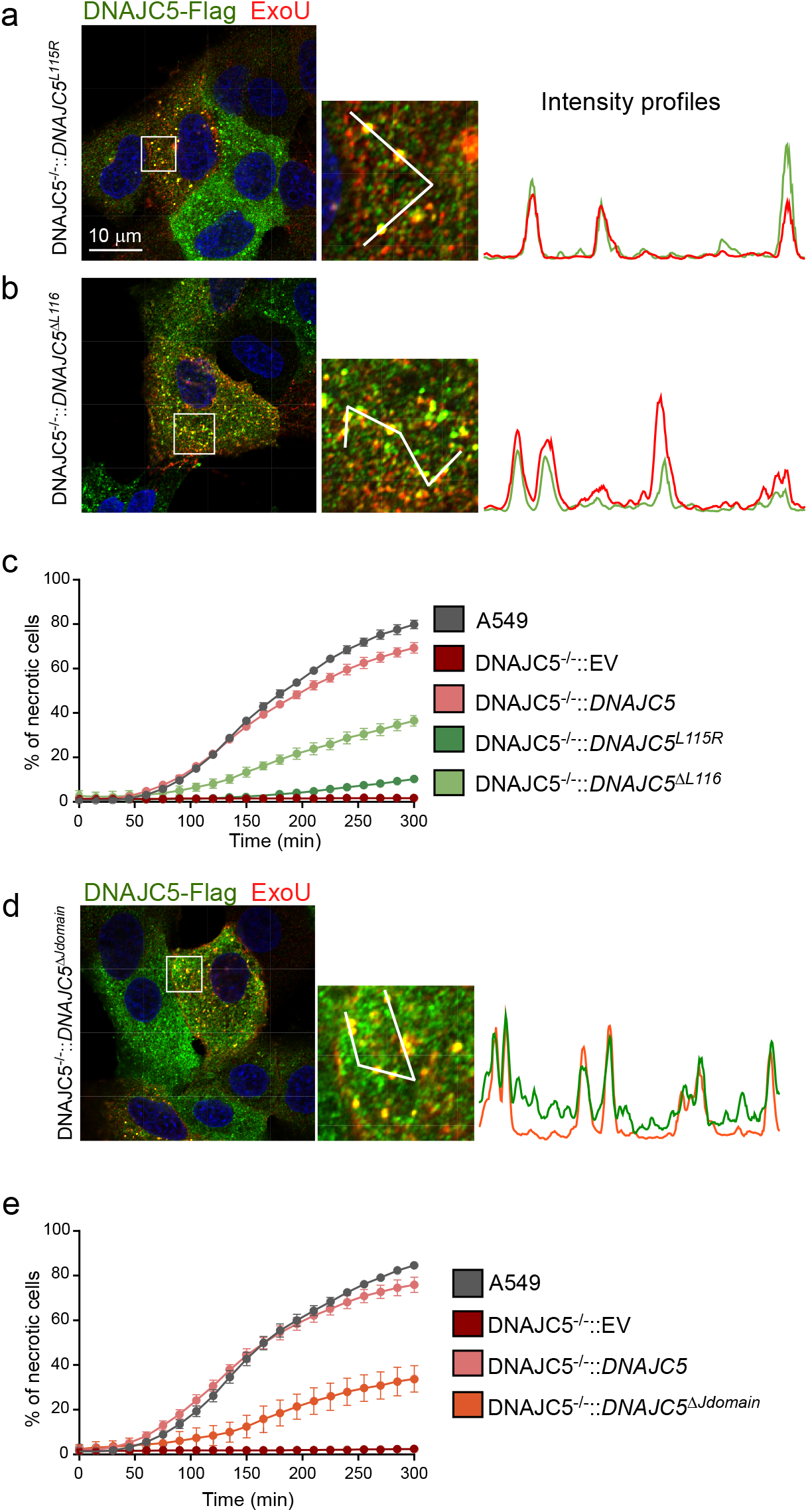
DNAJC5 escort of ExoU to the plasma membrane is required for toxicity. Localizations by immunofluorescence of ExoU (red) and DNAJC5-FLAG (green) in DNAJC5^−/−^::DNAJC5L115R cells infected with CHA-*exoUS142A* on a z-section. A selected area was enlarged and a bar was drawn, to establish the intensity profiles for both signals. **b**. Immunofluorescence localizations of ExoU (red) and DNAJC5-FLAG (green) in DNAJC5^−/−^::DNAJC5ΔL116 cells infected with CHA-*exoUS142A*. Intensity profiles were obtained as above. **c**. Cytotoxicity assay with DNAJC5^−/−^::DNAJC5L115R and DNAJC5^−/−^::DNAJC5ΔL116 cells, alongside controls, infected with PA14. N = 6 fields per condition. **d**. Immunofluorescence localizations of ExoU (red) and DNAJC5 (green) in DNAJC5^−/−^::DNAJC5ΔJdomain cells infected with CHA-*exoUS142A*. Intensity profiles were obtained as above. **e**. Effect of J domain deletion. Control cells and DNAJC5^−/−^ cells complemented with full-length DNAJC5, DNAJC5ΔJdomain or EV were infected with PA14, and cytotoxicity was recorded. (n = 6).

To confirm these results, we produced another DNAJC5 mutant lacking the J domain, also forming protein oligomers^52^, which has been hypothesized to alter DNAJC5 transportation function. DNAJC5^ΔJdomain^ similarly lead to restricted DNAJC5 localization in the perinuclear region, with some enlarged vesicles (Supplementary. 4 and 5), possibly due to protein accumulation in the lumen, where ExoU and Lamp2 colocalized (Fig. 5d). As with the pathological mutations L115R and ΔL116, ExoU toxicity was dramatically reduced in DNAJC5^−/−^::DNAJC5^ΔJdomain^ cells (Fig. 5e).

Based on these results, we concluded that DNAJC5 mutations affecting vesicle trafficking to the plasma membrane block ExoU-driven cell necrosis.

### ExoU phospholipase activity is independent of DNAJC5

To assess whether DNAJC5 contributes to the activation of ExoU enzymatic activity *per se*, we expressed and purified recombinant ExoU and examined its catalytic activity in a PLA2 assay, in the presence of membrane or soluble fractions prepared from uninfected DNAJC5^−/−^ or DNAJC5^−/−^::DNAJC5 cells. Only the membrane fractions enhanced ExoU phospholipase activity (Fig. 6), suggesting that a membranous component, probably PI(4,5)P2, activates ExoU catalytic activity, as previously reported^14^. Importantly, no significant difference was detected when ExoU was incubated with membrane fractions from either DNAJC5^−/−^ or DNAJC5^−/−^::DNAJC5 cells, showing that DNAJC5 is not directly involved in ExoU phospholipase activity.

**Figure 6.**
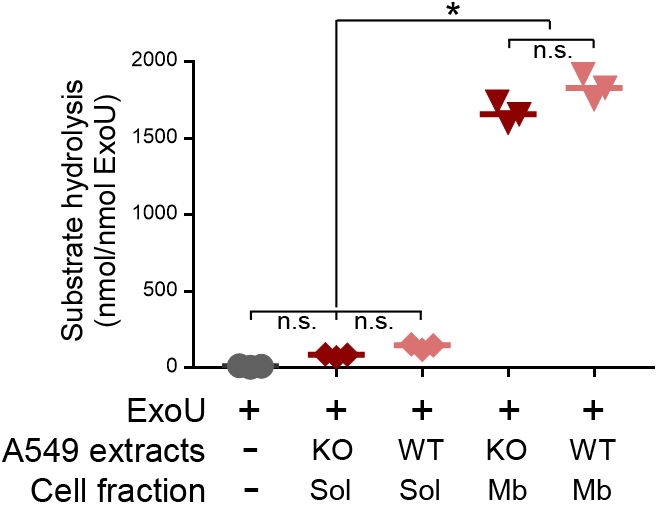
ExoU phospholipase activity is independent of DNAJC5. Phospholipase activity of purified ExoU (65 pmols) was measured in the presence of soluble (Sol) or membrane (Mb) fractions from uninfected DNAJC5^−/−^ (KO) or DNAJC5^−/−^::DNAJC5 (WT) cells. Experiments were performed in triplicates and incubated for 24 h. Data are expressed in nmoles of substrate hydrolyzed per nmoles of ExoU. Statistical differences were established by ANOVA (p < 0.0001), followed by Tukey’s test (*, p < 0.0001).

## Discussion

The aim of this study was to identify host factors required for full ExoU toxicity using a genome-wide screening approach. Our results demonstrate that the host chaperone DNAJC5 is required for the toxic activity of ExoU. In the bacterial cytoplasm, ExoU forms a complex with its cognate chaperone, SpcU, from which it dissociates prior to travel, probably unfolded, through the injectisome^53^. Once injected into the host cell, ExoU partly colocalizes with DNAJC5+ vesicles in the perinuclear zone (i.e., LEs) and at the cellular periphery.

DNAJC5 and Lamp2 only colocalized in perinuclear vesicles, and Lamp2 labelling was lost when DNAJC5+ vesicles were exported to the cellular periphery (Supplementary. 4). Similar patterns were reported in Cos7 cells with Lamp1, another late endosome/lysosome marker^29^, indicating that peripheral DNAJC5+ vesicles cannot be considered strictly “late endosomes”.

DNAJC5 has been linked to an unconventional secretion system (MAPS), both for the translocation of misfolded cytosolic proteins into the vesicle’s lumen and for the transport of these vesicles to the plasma membrane. DNAJC5-dependent protein exocytosis uses one of the two processes: either the vesicle fuses to the plasma membrane, allowing the release of the vesicle’s intraluminal content into the extracellular milieu^27^, or exosomes are produced^30^.

Unlike classical MAPS cargos, ExoU localized to the vesicle’s limiting membrane (Fig. 3d), it is therefore probably not translocated into the vesicle’s lumen, and remains at the external side. This position would allow ExoU to target the inner leaflet of the plasma membrane once the vesicle reaches the cell’s periphery (Supplementary Fig. 6). Moreover, MAPS cargos need Hsp70/Hsc70 chaperones for secretion through this pathway^26^, while ExoU transport is Hsp70/Hsc70-independent (Fig. 4). This lack of chaperone-dependence is probably linked to its position on the external side of the vesicle membrane.

ExoU toxicity was altered by mutations known to alter DNAJC5 function in vesicle trafficking, i.e., L115R and ΔL116^31,34,35,51^, as well as J domain deletion. These results confirm that ExoU, once delivered into the host’s cytoplasm, uses the DNAJC5-associated secretion machinery for transport to the plasma membrane. This result also suggests that patients with adult neuronal ceroid lipofuscinosis might be more resistant to ExoU intoxication.

Although ExoU and DNAJC5 colocalized at the vesicle’s limiting membrane, our attempts to identify a physical association between ExoU and DNAJC5 were unsuccessful. Furthermore, lack of DNAJC5 did not prevent ExoU from associating with membranes (Fig. 3a,b), confirming that ExoU binding to intracellular vesicles is DNAJC5-independent. Therefore, although DNAJC5 is an essential component of the ExoU transportation pathway, it is not the receptor for ExoU at the vesicle’s surface. Consequently, further studies will be required to identify the receptors allowing ExoU to dock to vesicle membranes.

The importance of ubiquitin binding to ExoU to stabilize the ExoU-membrane association and enhance its catalytic activity was demonstrated in previous studies^10,11,54,55^. It will now be interesting to examine whether ubiquitin binding also facilitates or strengthens the association between ExoU and DNAJC5+ vesicles, as it does at the plasma membrane.

Several bacterial toxins exploit vesicle trafficking for retrograde transport^56^, including T3SS cytotoxins^57,58^. In particular, *P. aeruginosa*’s ExoS toxin moves to the perinuclear region in a microtubule& and dynamin-dependent process, suggesting involvement of the endocytic pathway^59^. Interestingly, ExoU was observed to associate with DNAJC5-vesicles, particularly in the lower part of the cell (Fig. 3c). We previously showed that ExoU partially colocalized in cytoplasmic vesicles with EEA1^43^, a protein marker of early endosomes. It is thus possible that ExoU reaches the perinuclear region by an endocytic pathway like that exploited by ExoS, because vesicle transport is faster than free diffusion in the cytosol.

According to Deng et al.^59^, ExoS recycles to the plasma membrane by an unknown process. Our results showing full ExoS toxicity in DNAJC5^−/−^ cells (Supplementary Fig. 2a,b) demonstrate that ExoS and ExoU use distinct pathways to reach the plasma membrane.

As reported previously, only PI(4,5)P2, localized at the plasma membrane, can induce a conformational change in ExoU structure, leading to ExoU oligomerization and activating its catalytic activity^12,14,60^. In agreement with this specific activation mechanism, our results show that DNAJC5-dependent plasma membrane targeting is required for ExoU in cytoplasmic vesicles to induce membrane disruption (Fig. 1c).

In conclusion, we have elucidated a favored transportation system targeting ExoU to the only cellular location where it is active, i.e., the plasma membrane. Importantly, our initial screen for factors involved in this targeting identified only *DNAJC5*, suggesting that the protein it produces may be the Achilles’ heel for this highly potent toxin. Inhibitors of DNAJC5 (with a better efficiency than quercetin) or the MAPS pathway could be of considerable interest for adjunct therapy to treat infections with ExoU+ *P. aeruginosa* strains.

## Materials and Methods

### Lentiviral production using TKOv3 library and MOI determination

Toronto human knockout pooled library (TKOv3) was a gift from Jason Moffat and obtained from Addgene (#90294). It is a one-component library with guide-RNAs inserted in lentiCRISPRv2 backbone as well as the *cas9* gene. This library contains four gRNAs targeting each of the 18,053 protein coding genes and control gRNAs targeting EGFP, LacZ and luciferase (71,090 total gRNAs). The gRNA library-expressing lentiviruses were produced as described in Moffat’s lab protocol (REV.20170404). The library was amplified in Lucigen Endura electrocompetent cells (#60242) to reach at least 200x colonies per guide-RNA and the library plasmid pool was purified using NucleoBond Xtra-Maxi Kit (Macherey-Nagel, #740414.50). Then, lentiviruses were produced by transfection of HEK293T cells with the library plasmid pool. In brief, X-tremeGENE 9 DNA Transfection reagent (Roche, #06365787001) was diluted in Opti-MEM serum-free media. Following 5 min of incubation at room temperature, an appropriate mixture of plasmids was added in a 3:1 ratio of Transfection Reagent: DNA complex. This mixture of plasmids was composed of the library plasmid pool and the lentiviral packaging and envelope plasmids (psPAX2 and pMD2.G) at a 1:1:1 molar ratio (8 μg:4.8 μg:3.2 μg, respectively). The solution was then mixed and incubated at room temperature for 30 min. Thereafter, the transfection mix was added to 70-80% confluence HEK293T cells in a drop-wise manner and cells were incubated for 24 h. Following this incubation time, media were replaced by fresh DMEM, containing 6% of bovine serum albumin (BSA). The next day, the lentiviruses-containing media were harvested, centrifuged to pellet any packaging cells, and supernatants were stored at −80°C. The lentiviral concentration was established by serial dilutions on A549 cells, as described below, treated or not treated with 2 μg.mL^−1^ puromycin. The virus volume that gave 30% survival with puromycin selection vs without puromycin was chosen for library construction, in order to limit the number of lentiviruses per cell.

### Construction of CRISPR-Cas9 library in A549 cells and ExoU screen

The gRNA library-expressing A549 cells (hereinafter named A549-CRISPR cells) were constructed as described in Moffat’s lab protocol. Briefly, 5.10^7^ trypsinized A549 cells (calculated for 200-fold TKOv3 library coverage) were prepared in DMEM containing 10% FBS, supplemented with 8 μg.mL^−1^ polybrene to enhance lentiviral infection. Cells were incubated with lentiviral particles at MOI of 0.3. After 24 hours of incubation, media were replaced with DMEM containing 10% FBS supplemented with 2 μg.mL^−1^ puromycin and cells were additionally incubated for 48 h to select transfected cells. The library was frozen at −80°C before the screen.

For the screen, 15×10^6^ A549-CRISPR cells were plated on a 15-cm dish and infected with PA14 for 90 min at MOI of 10 to reach 20-30% of surviving cells. To stop the infection, cells were trypsinized after washing and reseeded in DMEM supplemented with 10% FBS and 20 μg.mL^−1^ polymixin B. The surviving population was expanded and evaluated daily to monitor the recovery of cells. The day after the seeding, 90 μg.mL^−1^ gentamicin was also added in the medium to prevent proliferation of polymyxin-resistant bacteria. When surviving cells reached 70-80% confluence, they were subjected to a second and a third round of infection with PA14, allowing the repetition of the procedure with the same library coverage. The screen on A549-CRISPR cells was done in biological triplicates.

### Genomic DNA extraction, sequencing and analysis

After three rounds of infection, the surviving populations were expanded to obtain 2.10^7^ cells. Uninfected A549-CRISPR cells were expanded similarly. Each cell population was subjected to genomic DNA (gDNA) extraction with a QIAamp DNA Blood Maxi kit (Qiagen, #51194) by following the manufacturer’s protocol. Then, a one-step PCR was carried out to enrich and amplify gRNAs from the genome of selected cell populations with Illumina TruSeq adapters (i5 and i7 indices). Moreover, a unique barcode of two 8-bp index required for Illumina sequencing was added during the one-step PCR to each pool of amplified gRNAs.

The PCR conditions for 50 μL were 25 μl of GoTaq G2 Hot Start Green Master Mix (#M7423, Promega), 0.5 μM of each primer (Supplementary Table 3) and 2.5 μg of gDNA. Three PCR reactions were performed simultaneously per sample to obtain sufficient quantities of amplified products. The PCR program was: denaturation at 95°C for 2 min, followed by 26 cycles at 95°C for 30 s, 55°C for 30 s and 65°C for 30 s, and a final elongation step at 65°C for 5 min. After PCR amplification, each 50-μL reaction from the same sample were pooled, electrophoresed and the 200-bp bands were excised from agarose gel slice using Monarch DNA Gel extraction Kit (NEB, #T1020S). Then, each sequencing library was quantified on both NanoDrop and Qubit and a quality control of DNA was performed on Agilent Bioanalyzer system. Finally, a high throughput sequencing was performed on the pool of amplicons using a NextSeq 500 device (Illumina) at the CNRS platform of Orsay (Institut de Biologie Intégrative de la Cellule) and raw data were processed and analysed using a web-based analysis platform named CRISPR-AnalyzeR. The adjusted p-values were analysed according to MAGeCK method to identify overrepresented genes targeted by gRNAs in output compared to input.

### Cytotoxicity assay

For infection assays, 1.5×10^4^ A549 cells or derivatives were seeded per well in a 96-well plate 48 h before infection in DMEM supplemented with 10% FBS and 200 μg.mL^−1^ neomycin for transfectants. Thirty minutes before infection, Syto24 was added to the medium at 0.5 μM to label cell nuclei. Then, medium was removed and replaced by DMEM supplemented with PI at 1 μM. Cells were subsequently infected at MOI 20 with bacteria (OD1) unless indicated. The kinetics of PI incorporation was followed using an IncuCyte live-Cell microscope (Sartorius). Acquisitions were done every 15 min for 5 h using a 10X objective. Images from bright field (phase), green channel (acquisition time 200 ms) and red channel (acquisition time 400 ms) were collected. The percentage of necrotic cells was calculated by dividing the number of PI-positive cells by the number of Syto24-positive cells.

For the cell retraction assay, cells were labelled with the CellTracker Red CMTPX (1 μM). Images were treated with ImageJ software. Briefly, images of CellTracker staining were binarized and total cell surface was calculated for 6 images at each time point.

### Immunofluorescence microscopy

For ExoU, DNAJC5, FLAG and Lamp2 stainings, 5×10^5^ A549 cells or derivatives were seeded in each well of a 24-well plate 48 h before fixation. For the co-staining with ExoU, cells were previously infected with CHA-*ExoU*^*S142A*^ or CHA-*ExoU*^S142AΔJdomain^ for 4 h at MOI of 10. Cells were fixed with 4% paraformaldehyde (PFA) for 15 min at room temperature and permeabilized with 0.5% Triton X-100 in PBS containing 4% PFA for 5 min. Cells were then stained using standard procedure with appropriate primary and secondary antibodies. Then, cells were counterstained with Hoechst. Images were collected on a Zeiss LSM880 confocal equipped with a Zeiss Plan-APO ×63 numerical aperture 1.4 oil immersion objective.

Successive planes in 3D stacks were taken every 0.2 μm. Images of z-sections were analysed using the Intensity profile module from Icy software.

### *Drosophila* mutant engineering and infection assay

To silence the *Csp* gene in *Drosophila melanogaster*, RNA interference was used. In brief, two different transgenic fly lines *Csp*-KD1 (Stock #34168 from the Vienna *Drosophila* Resource Center) and *Csp*-KD2 (Stock #33645 from the Bloomington *Drosophila* Stock Center) expressing a shRNA or a long dsRNA targeting *Csp*, respectively, under the control of GAL4-responsive elements were used. These flies were crossed with transgenic flies expressing the *Gal4* gene under the control of a temperature-inducible promoter (hs-Gal4, stock #2077 from the Bloomington *Drosophila* Stock Center). Flies expressing a siRNA targeting the firefly *Luciferase* gene (Stock #31603 from the Bloomington *Drosophila* Stock Center) were also crossed with the heat shock-Gal4 line and used as control. The progeny was reared until the flies were 7 to 10-day old and the *Gal4* gene expression was induced by three heat shocks of 1 h at 37°C, each performed during three consecutive days, in order to silence the *Csp* gene. Flies were then infected with a stationary phase culture of PP34 strain (or PP34Δ*exoU* strain), diluted at OD 1, by thoracic needle pricking, as previously described^42^, and their survival rates were monitored. To confirm the knockdown of CSP, 20 flies per condition were homogenized in RIPA lysis buffer with a Precellys 24 (Bertin instruments) using CK14 tubes containing ceramic beads at 5,000 rpm for four cycles of 30 s. Lysates were analysed by Western blot using the anti-*Drosophila* CSP antibody.

### Statistics

Statistics on genomic screening data are described above.

GraphPad 7.04 software was used for all other statistical analyses. For cytotoxicity studies or cell retraction assay, no statistical test was used. For Bla activity assay, a Student’s t-test was used between conditions using ExoU-bla. For PLA2 activity test, a one-way ANOVA was employed, followed by Tukey’s post-hoc test for data comparison. For the two latter assays, data distribution was normal according to Shapiros-Wilk’s test. For fly infection, a Log-Rank test was used. Data were considered significantly different when p < 0.01.

## Supporting information

Supplemental methods, figures, Tables and References

## Data availability

The sequencing data have been deposited in the NCBI Gene Expression Omnibus (GEO) and are accessible through GEO accession number GSE154751.

## Acknowledgments

We are grateful to Luke Chamberlain for the gift of the EGFP-DNAJC5 expression plasmids, to Simona Barzu for the ExoU antibody, Michel Ragno for ExoU purification. Confocal microscopy was performed at the μLife platform and automated microscopy at the CMBA platform. International Health Management Association (IHMA, USA) kindly provided the *P. aeruginosa* IHMA879472 strain. We thank Laurence Aubry and Agnès Journet for helpful discussions.

## Author contribution

V.D. and S.B. performed most experiments and analyzed data. P.H. and V.J. performed some experiments. E.T. and M-O.F conceptualized the *Drosophila* experiments and managed the fly facility. I.A. provided the bacterial models. P.H. conceptualized the project and wrote the manuscript.

## Supplementary information

Supplementary methods, figures and tables, and additional references, are provided as supplementary information.

