## Supplemental methods, figures, Tables and References for "The bacterial toxin ExoU requires a host trafficking chaperone for transportation and to induce necrosis"

### Supplementary Methods

#### Bacterial strains and plasmids

The bacterial strains and plasmids used in this study are listed in Supplementary Table 1. Bacterial strains were grown in Luria-Bertani (LB) medium at 37°C with vigorous shaking (300 rpm). When mentioned, antibiotics were added at the following concentrations: 100 µg.mL$^{-1}$ ampicillin and 300 µg.mL$^{-1}$ carbenicillin. For infection assays, overnight cultures were diluted to optical density ($OD_{600}$) of 0.1 and grown under agitation to reach $OD_{600}$ of 1.

For construction of PP34Δ*exoU*, the internal nucleotide sequence of *exoU* from PP34 isolate[61] was amplified by PCR and cloned into pEX100T[62]. The gentamicin cassette was extracted from pUC-Gm and inserted into the unique *Eco*RI site of *exoU*. Then, the *exoU*-Gm fragment was introduced into the genome of *P. aeruginosa* PP34 by allelic exchange following standard triparental mating and *sacB* selection technique. The mutant was verified by PCR using primers ExoU-Gm_Fw and ExoU-Gm_Rev (Supplementary Table 2) and resequencing.

For *exoU-bla* fusion, the DNA fragment encompassing the *exoU* promotor and coding sequence was amplified by primers ExoU-BamHI and ExoU-XbaI (Supplementary Table 2) using pIA*exoU*$^{S142A}$*spcU* as template[43]. The PCR fragment was then transferred into *Bam*HI- and *Xba*I-digested pIA*exoS-bla* plasmids, where *exoS* was excised. The expression and secretion of ExoU-Bla was verified by immuno-blotting using anti-ExoU antibodies. The plasmids were introduced into PP34Δ*exoU* by transformation[63].

#### Purification of His6-ExoU

The *exoU* gene amplified from the PP34 isolate (GESPA collection[61]) was cloned in pET15b. The obtained plasmid pET15b-*exoU* was introduced into *E. coli* BL21 (DE3) Star (InVitrogen). The induction was obtained by adding IPTG at 1mM concentration during 3 h. The bacterial pellet, resuspended in 25 mM Tris-HCl, pH8, 500 mM NaCl and 10 mM imidazol, was disrupted using a Microfluidizer. Purification of $His_6$-ExoU was performed on AKTA purifier using a HiTrap HP 5-ml column (GE HealthCare) with a step gradient of imidazole. Fractions, eluted with 200 mM imidazole and containing the ExoU protein, were loaded onto HiLoad SD200 16/60 prep column for a second step of purification.

#### Antibodies and reagents

The rabbit polyclonal antibody targeting human CSPα (DNAJC5) was purchased from ThermoFisher (#PA1-776). The mouse polyclonal antibodies against β-actin (#A1978), β-tubulin (#T0198) and FLAG (#F1804) were purchased from Sigma-Aldrich. Specific antisera for ExoU were obtained in rabbits with 50 µg of purified full length recombinant ExoU. The specific antibodies were affinity-purified on $His_6$-ExoU. The mouse monoclonal antibody targeting CSP in *Drosophila melanogaster* (named DCSP-1) was purchased from DSBH (#ab49). DCSP-1 (ab49) was deposited to the DSHB by Buchner, E. / Hofbauer, A. The monoclonal mouse Hsc70 (HSPA8) and Hsp70 (HSPA1A) antibodies were purchased from R&D Systems (#MAB4148 and #MAB1663-SP, respectively). The anti-Lamp2 antibody was from BD Transduction Laboratories (#555803). Neomycin (#10131-035), Syto24 (#S7572) and propidium iodide (PI)(#P3566) were purchased from GIBCO, ThermoFisher and InVitrogen, respectively. The CellTracker Red CMTPX (#C34552) and CCF2-AM (K1032) were from ThermoFisher. Quercetin (#Q4951) and probenecid (#P8761) were from Sigma-Aldrich. siRNAs against Hsp70 (HSPA1, #s6968) and Hsc70 (HSPA8, #s6985) were purchased from

ThermoFisher. The Complete inhibitor cocktail was from Roche (#04693159001) and the Micro-BCA Protein Assay Kit from ThermoFisher (#23235). The Lipofectamine RNAiMax kit was from Invitrogen (#13778-075).

**Eukaryotic cell lines and growth conditions**
The human embryonic kidney (HEK) 293T and A549 cells and their derivatives were cultured in Dulbecco's modified Eagle's medium (DMEM) supplemented with 10% foetal bovine serum (FBS). Cells were grown at 37°C with 5% $CO_2$ and routinely passaged when reaching 70 to 80% confluence. Transfected cells with pLVX-*IRES-neo* vector, containing the wild-type or mutated *DNAJC5-FLAG* gene, were maintained by addition of 200 $\mu g.mL^{-1}$ neomycin to the supplemented medium.

**Knockout of *DNAJC5* gene**
A pair of oligonucleotides corresponding to the enriched gRNA targeting *DNAJC5* gene (Supplementary Table 4) were annealed and cloned into the *BsmBI*-digested pLentiCRISPRv2 vector. Then, as previously described for TKOv3 library, lentiviral particles containing the recombinant plasmid pLentiCRISPRv2-gRNA-*DNAJC5* were created using the HEK293T cells and the psPAX2 and pMD2.G plasmids. Lentiviruses containing the pLentiCRISPRv2 vector without gRNA were also produced and referred to as empty vector (EV). A549 cells were then infected with these lentiviruses using 2 mL of lentiviral particles per 10-cm dish with cells at 50% confluence. Transfected cells were selected with 2 $\mu g.mL^{-1}$ puromycin for 48 h and clones were isolated by limiting dilution. Clones were selected for their absence of DNAJC5 expression by Western blot and immunofluorescence. One clone was selected for further experiments.

**Complementation of deficient cells with *DNAJC5-FLAG* gene and its derivatives DNAJC5$^{H43Q}$ and DNAJC5$^{S10A-S34A}$**
The *DNAJC5* wild-type gene was synthetized by the Genewiz company and cloned into pUC57 plasmid. Some modifications have been introduced, without changing the amino acid sequence, to prevent the Cas9 endonuclease from cleaving the gene when inserted into the genome of deficient cells. Moreover, the 3X-FLAG tag was designed upstream of the gene and *Eco*RI and *Bam*HI restriction sites were added upstream and downstream of the *DNAJC5-FLAG* gene respectively (Supplementary Table 5). The gene was inserted into pLVX-IRES-neo vector in *Bam*HI and *Eco*RI sites. Lentiviral particles containing the empty vector pLVX or the pLVX-*DNAJC5-FLAG* plasmid were produced as previously described. Thereafter, $2.6x10^6$ DNAJC5$^{-/-}$ cells were seeded per 10-cm dish the day before and were infected with 2 mL of lentivirus for 24 h. Then, media were replaced and transfected cells were cultured in DMEM containing 10% FBS and supplemented with 800 $\mu g.mL^{-1}$ neomycin for at least 8 days before decreasing the antibiotic concentration to 200 $\mu g.mL^{-1}$. In the same way, H43Q and S10A-S34A mutations on *DNAJC5-FLAG* gene were introduced by synthetizing the mutated genes (Supplementary Table 5) and by infecting the deficient cells with lentiviral particles containing the mutated genes cloned into the pLVX vector. In each case, clones were isolated by limiting dilution and tested for DNAJC5 expression by Western blot and immunofluorescence.

**Site-directed mutagenesis of *DNAJC5***
The QuikChange Site-Directed Mutagenesis Kit (Agilent, #200523) was used to perform mutations on *DNAJC5-FLAG* gene. Briefly, oligonucleotide primers containing the desired

mutations flanked by unmodified nucleotide sequence were synthetized as recommended by the manufacturer guidelines (Supplementary Table 6). For mutagenesis, 125 ng of each of the two complementary oligonucleotides were used in a reaction volume of 50 µL containing 30 ng of pUC57-*DNAJC5*-FLAG plasmid, 1 µL of dNTP mix and 5 µL of 10X reaction buffer. Then 1 µL of *PfuTurbo* DNA polymerase at 2.5 $U.\mu L^{-1}$ was added and the mixture was PCR amplified using the following cycling parameters: 95°C for 30 s followed by 25 cycles at 95°C for 30 s, 55°C for 1 min and 68°C for 3 min, and a final elongation step at 68°C for 5 min. Thereafter, the nonmutated plasmid was digested by adding 1 µL of DpnI at 10 $U.\mu L^{-1}$ directly to amplification reaction followed by an incubation at 37°C for 1 h. The remaining plasmid containing the desired mutation was transformed into Quick Change XL1-Blue Supercompetent cells. Mutants were checked by DNA sequencing. Then, as previously described, the mutated *DNAJC5* gene was digested and cloned into the pLVX-IRES-neo vector and deficient cells were complemented with the mutated gene. Clones were isolated by limiting dilution and selected as above.

**Phospholipase activity assay of ExoU**

The phospholipase activity of ExoU was performed, as reported previously[13], using the Cayman Chemical $cPLA_2$ kit (#765021). Briefly, 5 µL of purified 6His-ExoU protein at 1 $mg.mL^{-1}$ (65 pmols) were used per well of a 96-well plate containing 5 µL of Assay Buffer and 5 µL of cytosolic or membrane fractions (normalized by $A_{280}$) from DNAJC5$^{-/-}$ cells and DNAJC5$^{-/-}$::DNAJC5 cells. Then, reactions were initiated by adding 200 µL of substrate solution containing 1.5 mM arachidonyl thiophosphatidylcholonie and by shaking the plate for 30 s followed by an incubation for 1 h at room temperature. Absorbance was monitored at 24 h at 405 nm with an automated plate reader (Spark 10M by TECAN) after addition of 10 µL of a solution containing 25 mM 5,5-dithiobis (2-dibitrobenzoic acid) (DTNB). The $PLA_2$ activity of ExoU was also monitored in wells without extracts called "Blank wells". Experiments were performed in triplicate. The $PLA_2$ activity of ExoU was calculated with the following formula where 10 is the extinction coefficient for DTNB.

$$\text{Substrate hydrolysis} = \frac{[\text{Mean of Abs (sample)} - \text{Mean of Abs (blank)}]}{10 \text{ x } 65.10^{-3} (nmol\ of\ ExoU)}$$

**ExoU injection assay**

A549 or DNAJC5$^{-/-}$ cells were seeded in 96-well plates 2 days before infection at $1.5 \times 10^4$ cells per well. Cells were infected at MOI of 10 for 4 h with the PP34Δ*exoU* strain producing ExoU$^{S142A}$-Bla toxin, as reported previously[36]. Then, cells were washed with PBS containing 2.5 mM probenecid, and incubated with freshly prepared CCF2-AM solution (2 µM) for 90 min in the dark at room temperature. The CCF2-AM cleavage by ExoU$^{S142A}$-Bla translocation was measured by comparing the emitted fluorescence at 447 nm (green, uncleaved) and 530 nm (blue, cleaved) upon excitation at 405 nm.

***DNAJC5* sequence analysis in DNAJC5$^{-/-}$ cells**

Genomic DNA was isolated from A549 and DNAJC5$^{-/-}$ cells ($10^7$ cells) using the QIAamp DNA Blood Maxi kit (Qiagen, #51194) and by following manufacturer's protocol. To amplify the *DNAJC5* gene, PCR reactions were carried out with 2.5 µg of gDNA in GoTaq G2 Hot Start Green Master Mix (Promega, #M7423) containing 0.5 µM of primers DNAJC5-Fw and DNAJC5-Rev (see Supplementary Table 2). PCR conditions were: 2 min at 95°C, followed by 30 s at 95°C, 30 s at 56°C and 1 min at 72°C (26x), and 5 min at 72°C. The PCR products were Sanger-sequenced.

**Western blot analysis**
Cells were washed twice with cold PBS and lysed with lysis buffer containing 1% Triton X-100, 50 mM Tris-HCl pH 7.4, 150 mM NaCl, 1 mM EDTA, Roche protease inhibitor cocktail as well as vanadate (1 mM) and okadaic acid (50 nM). The lysates were centrifuged at 18,400 g for 10 min at 4°C and protein concentration in the supernatant was measured with the Micro-BCA Protein Assay Kit. Supernatants were then denatured using Laemmli buffer containing β-mercaptoethanol at 95°C for 5 min. For gel electrophoresis, proteins were run using 10% or 4-12% polyacrylamide gels (BioRad, #3450123). Proteins were then transferred onto a PVDF membrane, using the BioRad semi-dry transfer apparatus, and incubated for 1 h with 5% non-fat dairy milk followed by overnight incubation at 4°C with primary antibodies. Membranes were then probed with secondary HRP-antibodies for 90 min at room temperature. After washings, signals were detected using the Immobilon Western blot Substrate and the ChemiDoc MP Imaging System.

**siRNA transfection**
A549 cells were seeded at $1.5 \times 10^5$ cells per well in a 6-well plate (for cell lysate preparation) or at $1.5 \times 10^4$ cells per well in a 96-well plate (for cytotoxicity assay) 24 h prior to transfection. Transfection was carried out with the 3 pmols siRNAs per well of 96-well plate or 30 pmols per well of 6-well plates, using the Lipofectamine RNAiMax kit according to the manufacturer's protocol. After 24 h of incubation, the media were replaced by DMEM containing 10% FBS. Experiments were performed at 48 h post-transfection.

**Quercetin assays**
For infection assays, A549 cells were seeded as previously described in 96-well plate, 48 h before. Quercetin was freshly prepared at 100 mM in DMSO. The day of infection, Syto24 was added to the medium at 0.5 µM 2 h before infection, the medium was removed and replaced by DMEM supplemented with PI and quercetin at the indicated concentrations. Cells were then infected with bacteria at OD 1 and at MOI 10. The kinetics of PI incorporation was followed as previously described using an automatized microscope IncuCyte.

**Cellular fractionation**
Cellular fractionation was performed as previously described[29], with some modifications. Briefly, infected or uninfected cells ($2.10^7$) were washed with ice-cold PBS and scraped in 800 µL of buffer A (10 mM Tris-HCl pH 7.4, 10 mM KCl, 2 mM MgCl, 1 mM DTT, with protease inhibitor cocktail). The cellular suspension was supplemented to 250 mM sucrose to prevent subcellular organelle damage. Cells were fragmented by passing 20 times through a ball bearing homogenizer (8.020-mm bore, EMBL, Heidelberg, Germany) with an 8.006-mm ball bearing. The cellular homogenate was centrifuged at 1,000 × g for 10 min to remove nuclei and unbroken cells. The supernatant was then ultracentrifuged at 100,000 x g for 30 min to sediment total microsome membranes. The pellet was resuspended in buffer A with sucrose, and was conserved at -20°C, as well as the soluble fraction. Samples were normalized with $OD_{280}$.

**Expression of DNAJC5-EGFP in HUVECs**
Human umbilical vein endothelial cells (HUVECs) were isolated according to previously described protocols[64]. Cells were cultured in endothelial-basal medium 2 (EBM-2; Lonza)

supplemented as recommended by the manufacturer. HUVECs were transfected using nucleofection (Amaxa, Lonza) according to the manufacturer's protocol. Briefly, the day before transfection, cells were seeded at a density of 30,000 cells/cm2 in EBM-2 medium supplemented with EGM-2 SingleQuots. Cells ($2\times10^6$) were pelleted by centrifugation (6 min at 1,000 rpm) prior to being resuspended in 100 µL of Nucleofector® solution, mixed with 2.5 µg DNAJC5-EGFP plasmid[51]. The cells were nucleofected by using the program A-034. DNAJC5-EGFP-transfected cells were used at 24 h post-transfection.

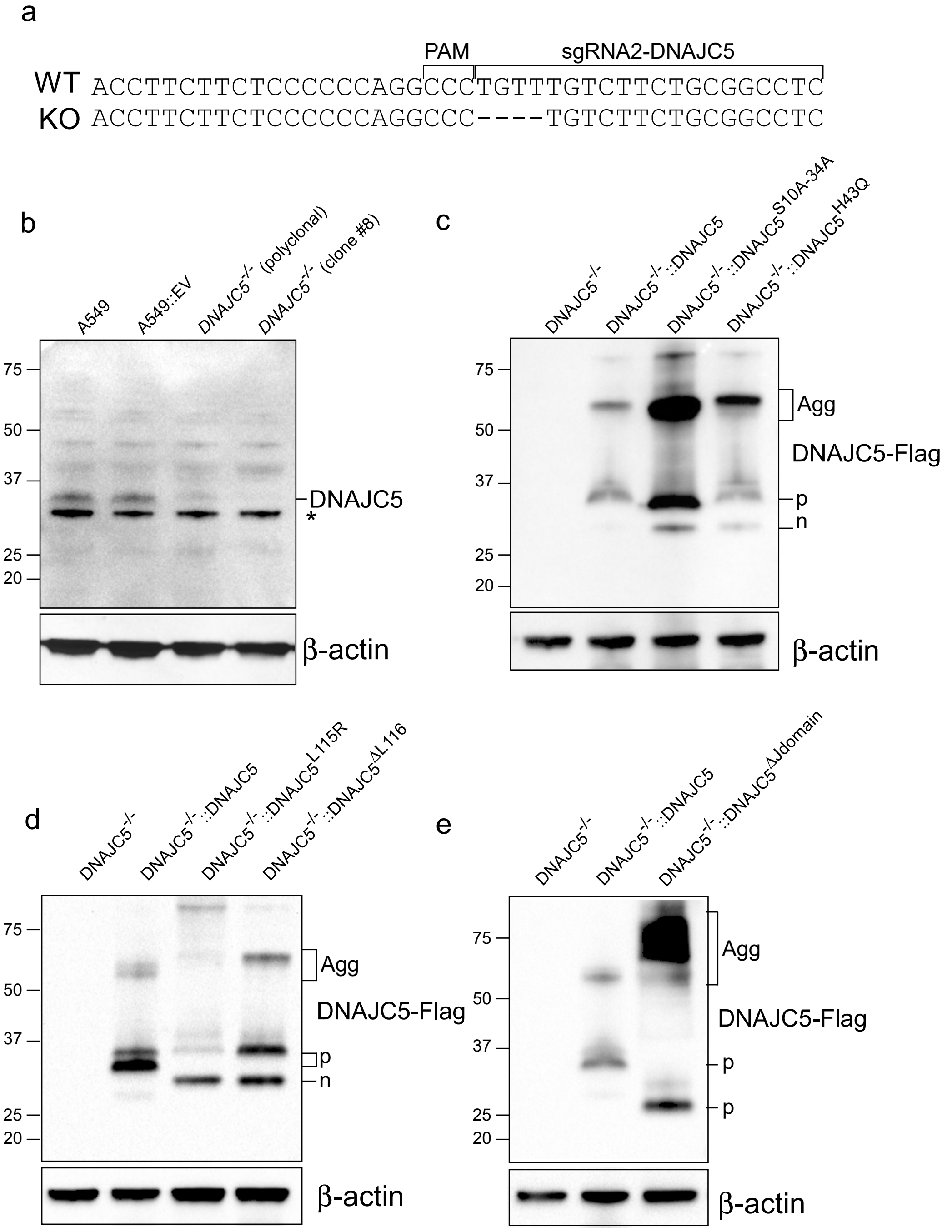


**Supplementary Fig. 1. Sequence of DNAJC5- gene clone #8 and expression of wild-type and mutant DNAJC5.**

**a**. Part of the native DNAJC5 sequence (WT), aligned with the mutated sequence present in the selected DNAJC5$^{-/-}$ cells clone #8. The four missing nucleotides are indicated. The same deletion was present on both alleles. The PAM sequence and the sequence of the gRNA used for inactivation are shown. **b**. Western blot with cellular extracts from A549, A549::EV and DNAJC5$^{-/-}$ (polyclonal population and clone#8) cells, revealed with DNAJC5 antibodies. ß-actin was used as loading control. **c**. Western blot showing DNAJC5-FLAG expression in DNAJC5$^{-/-}$::DNAJC5, DNAJC5$^{-/-}$::DNAJC5$^{H43Q}$ and DNAJC5$^{-/-}$::DNAJC5$^{S10A\text{-}S34A}$ cells, revealed with the FLAG antibodies. **d**. Western blot showing DNAJC5-FLAG expression in DNAJC5$^{-/-}$::DNAJC5, DNAJC5$^{-/-}$::DNAJC5$^{L115R}$ and DNAJC5$^{-/-}$::DNAJC5$^{\Delta L116}$ cells, revealed with the FLAG antibodies. **e**. Western blot showing DNAJC5-FLAG expression in DNAJC5$^{-/-}$::DNAJC5, DNAJC5$^{-/-}$::DNAJC5$^{\Delta Jdomain}$ cells, revealed with FLAG antibodies. Abbreviations: (n) native and (p) palmitoylated forms of DNAJC5; Agg: aggregates. (*) represents a non-specific band only visible in highly exposed images.

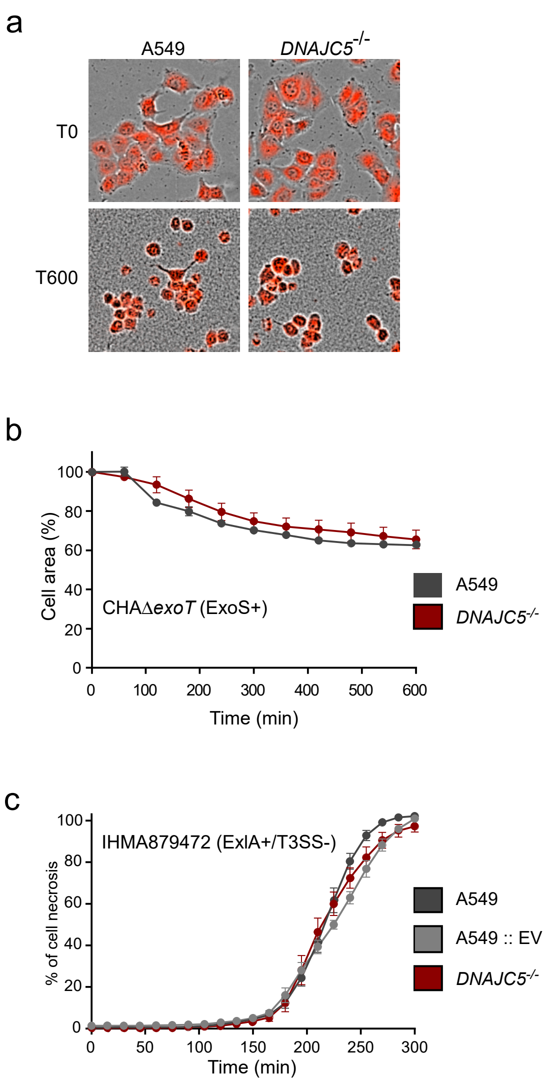


**Supplementary Fig. 2. DNAJC5 is not required for ExoS or ExlA toxicity.**

**a**. Intoxication with ExoS. A549 cells or DNAJC5$^{-/-}$ cells loaded with CellTracker Red CMTPX were infected with the CHAΔ*exoT* strain expressing ExoS. Intoxication was followed by time-lapse microscopy (n = 6). Images show merged acquisitions of phase contrast and CellTracker fluorescence at the beginning (T0) and at 10 hpi (T600). Both cell types displayed ExoS-dependent cell rounding. **b**. Quantification of ExoS-induced cell rounding using ImageJ software. **c**. A549, A549 ::EV and DNAJC5$^{-/-}$ cells were infected with the ExlA+ IHMA879472 strain, and cell necrosis was monitored based on PI incorporation (n = 6). EV, empty vector.

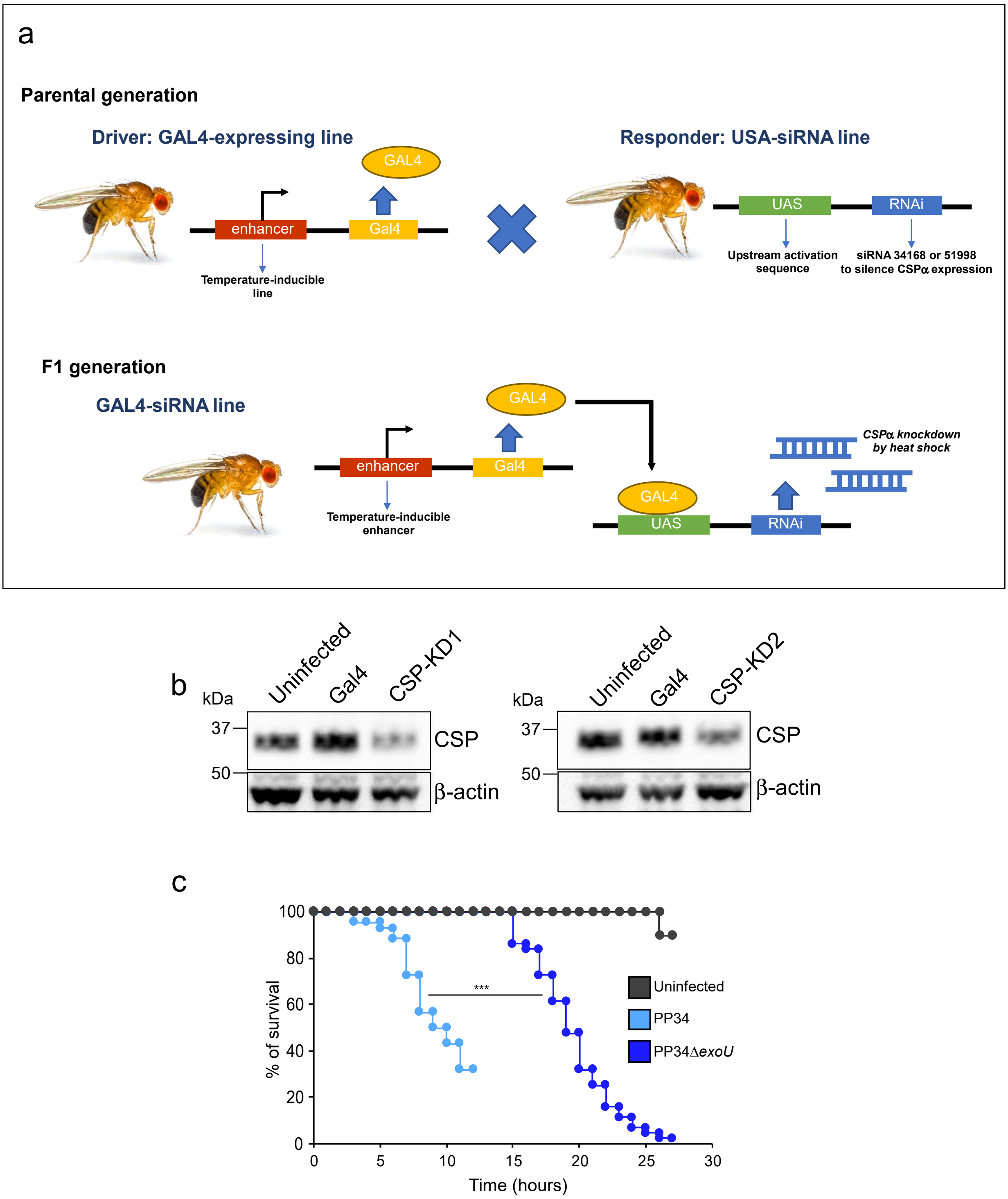


**Supplementary Fig. 3. Production of Csp knockdown in Drosophila and intoxication assay.** **a**. Mating scheme to generate *Csp*-KD flies. **b**. Western blots showing CSP levels in flies using two different RNAi transgenes (*Csp*-KD1 and *Csp*-KD2). ß-actin was used as loading control. **c**. Post-infection survival assay. Flies were infected by pricking the thorax with a needle dipped in suspensions of either PP34 (ExoU+; n = 50) or PP34Δ*exoU* (n = 50) strains. Data for mock-infected flies, pricked with PBS (uninfected; n = 10), are also shown. Statistical differences were determined using the Log-Rank test. ***: p <0.001.

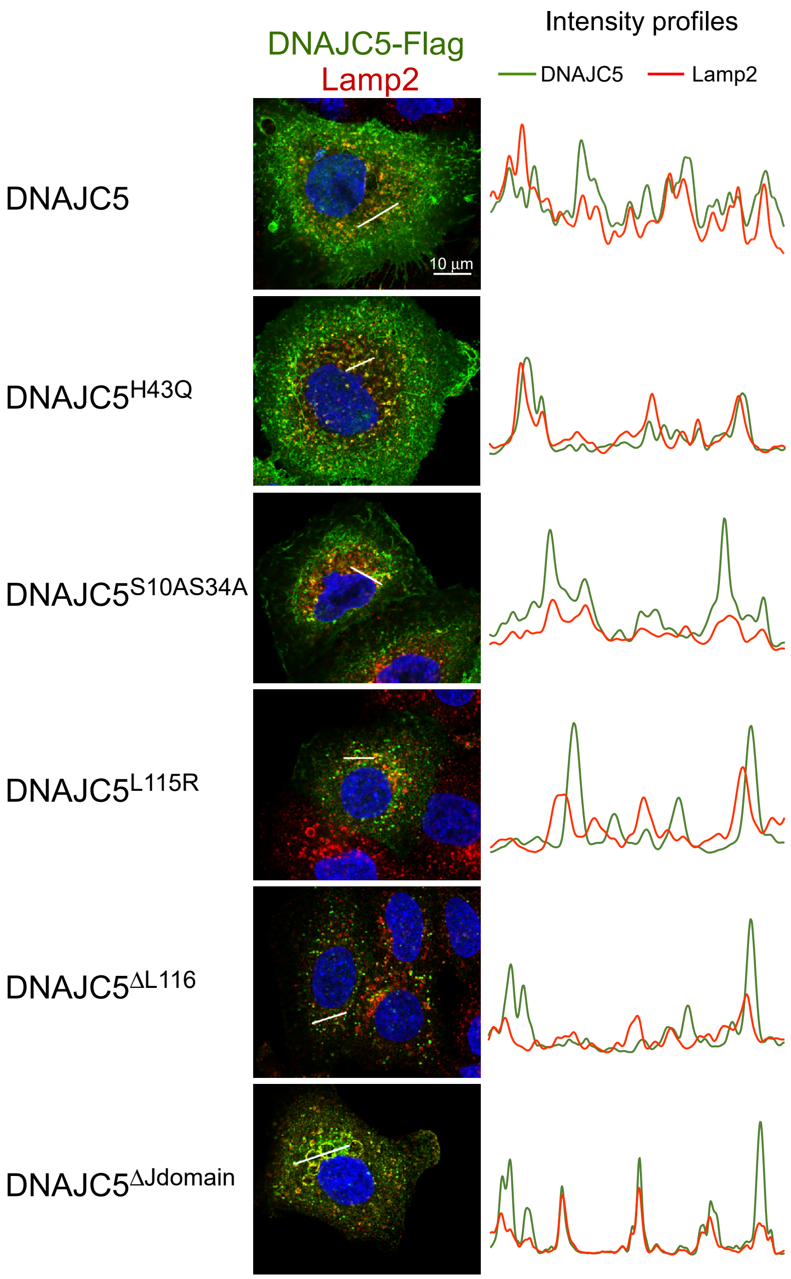


**Supplementary Fig. 4. Partial colocalization of DNAJC5 (wild type or mutants) with Lamp2.**

Immunostaining to reveal colocalization of DNAJC5 (green) and Lamp2 (red) in DNAJC5$^{-/-}$ cells complemented with DNAJC5-FLAG or a mutated form of DNAJC5, as indicated. Bars indicate where the intensity profiles for both labels were measured. Intensity profiles are shown on the right.

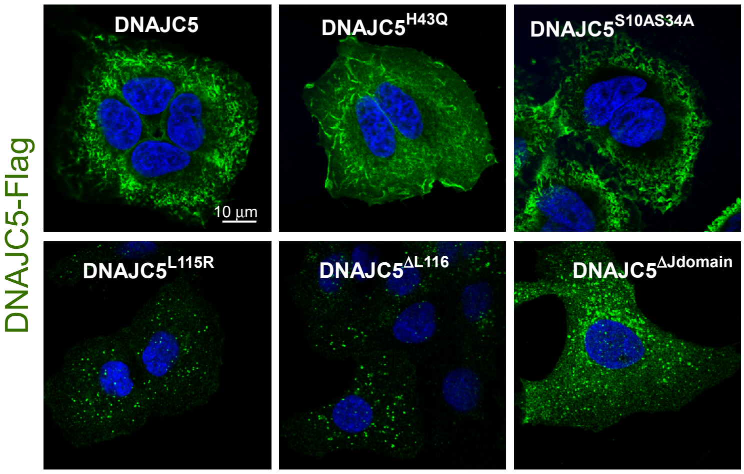


**Supplementary Fig. 5. Expression and localization of DNAJC5 mutants.** DNAJC5 and derivatives, fused to a FLAG tag, were expressed in DNAJC5$^{-/-}$ cells. Cells were fixed and labelled with an anti-FLAG antibody. One z-section image is shown for each cell line.

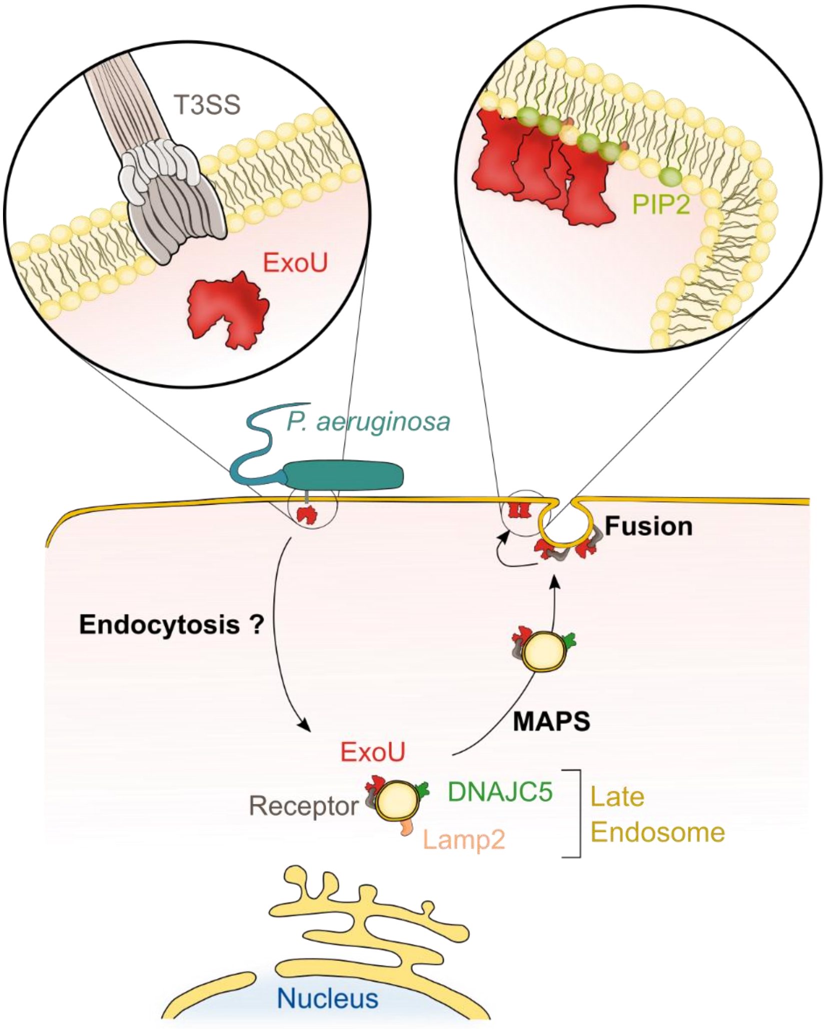


**Supplementary Fig. 6: Model of ExoU trafficking in host cells.** Upon delivery into the host cytoplasm by the T3SS, the toxin uses an endocytic pathway to reach the perinuclear region, as suggested by ExoU/EEA1 colabeling. Then, ExoU binds to the late endosome's limiting membrane (decorated by Lamp2 and DNAJC5), thanks to the interaction of ExoU with an as yet unidentified specific receptor at the vesicle's surface. ExoU remains at the external side of the vesicle's membrane and coopts the DNAJC5-dependent MAPS pathway to achieve anterograde transport toward the cellular periphery (where vesicles loose Lamp2), and eventually the plasma membrane (PM). Fusion of vesicles with the PM brings ExoU close to PM's inner leaflet, where its membrane localization domain binds to PI(4,5)P2. PI(4,5)P2 binding triggers conformational changes in ExoU, leading to toxin oligomerization and activation of its phospholipase activity, which eventually induces PM rupture.

**Supplementary Table 1: Strains and plasmids**

| Names | Relevant characteristics | Reference/Source |
|---|---|---|
| ***E. coli* strains** | | |
| One Shot™ TOP10 Chemically Competent *E. coli* | F- *mcrA* Δ( *mrr-hsd*RMS-*mcr*BC) Φ80*lac*ZΔM15 Δ *lac*X74 *rec*A1 *ara*D139 Δ( *araleu*)7697 *gal*U *gal*K *rps*L (StrR) *end*A1 *nup*G | Invitrogen, C404003 |
| NEB® 5-alpha Competent *E. coli* (High Efficiency) | *fhuA2 Δ(argF-lacZ)U169 phoA glnV44 Φ80 Δ(lacZ)M15 gyrA96 recA1 relA1 endA1 thi-1 hsdR17* | NEB, C2987H |
| Lucigen Endura electrocompetent cells | *recA13 supE44 ara-14 galK2 lacY1 proA2 rpsL20(StrR) xyl-5 λ– leu mtl-1 F– mcrB mrr hsdS20(rB–, mB–)* | Biosearch Technologies, 60242 |
| Quick Change XL1-Blue Supercompetent cells | *recA1 endA1 gyrA96 thi-1 hsdR17 supE44 relA1 lac [F´ proAB lacIqZΔM15 Tn10]* | Agilent, 200519-4 |
| Stbl3™ Chemically Competent *E. coli* | F-*mcr*B *mrrhsd*S20(rB-, mB-) *rec*A13 *sup*E44 *ara*-14 *gal*K2 *lac*Y1 *pro*A2 *rps*L20(StrR) *xyl*-5 λ-*leumtl*-1 | Invitrogen, C737303 |
| ***P. aeruginosa* strains** | | |
| PA14 | Contains *exoU* | [2] |
| PP34 | Contains *exoU* | [3] |
| IHMA879472 | T3SS⁻ and contains ExlB-ExlA two-partner system | [4] |
| PP34Δ*exoU* | ExoU⁻ | This study |
| PP34Δ*exoU::exoU-bla* | PP34Δ*exoU* strain expressing ExoU$^{S142A}$-Bla toxin | This study |
| CHAΔ*exoSexoT* | ExoS⁻ and ExoT⁻ | [5] |
| CHAΔ*exoSexoT* ::*exoU*$^{S142A}$(CHA-*exoU*$^{S142A}$) | ExoS⁻ and ExoT⁻ and ExoU⁺ catalytically inactivated; Cb$^R$ | [6] |
| CHAΔ*exoT* | ExoS⁻ and ExoT⁻ and ExoS⁺ | [5] |
| **Bacterial vectors** | | |
| pIA*exoS-bla* | Contains *exoS* gene downstream of the β-lactamase gene ; Cb$^R$ | [7] |
| pIA*exoU*$^{S142A}$*-bla* | pIA*exoS-bla* plasmid where *exoS* gene is excised and replaced by *exoU*$^{S142A}$; Cb$^R$ | This study |
| pUCP*ExoU*$^{S142A}$*spcU* | Contains *exoU* gene which is catalytically inactivated and *spcU* gene ; Cb$^R$ | [6] |

| | | |
|---|---|---|
| pUC57-*DNAJC5-FLAG* | Contains *DNAJC5* gene with N-terminal FLAG-tag; Amp$^{R}$ | This study |
| pUC57-*DNAJC5*$^{S10A\text{-}S34A}$-*FLAG* | Contains FLAG-tagged *DNAJC5* gene mutated on phosphorylation sites ; Amp$^{R}$ | This study |
| pUC57-*DNAJC5*$^{H43Q}$-*FLAG* | Contains FLAG-tagged *DNAJC5* gene mutated on the Hsc70/Hsp70 binding site; Amp$^{R}$ | This study |
| pUC57-*DNAJC5*$^{L115R}$-*FLAG* | Contains FLAG-tagged *DNAJC5* gene with L115R mutation in the cysteine-string domain ; Amp$^{R}$ | This study |
| pUC57-*DNAJC5*$^{\Delta L116}$-*FLAG* | Contains FLAG-tagged *DNAJC5* gene with the deletion of L116 in the cysteine-string domain ; Amp$^{R}$ | This study |
| pUC57-*DNAJC5*$^{\Delta Jdomain}$-*FLAG* | Contains FLAG-tagged *DNAJC5* gene with the deletion of the J-domain ; Amp$^{R}$ | This study |

**Mammalian vectors**

| | | |
|---|---|---|
| psPAX2 | Lentiviral packaging plasmid | Addgene, 12260 |
| pMD2.G | VSV-G envelope expressing plasmid | Addgene, 12259 |
| pLentiCRISPRv2 | Lentiviral vector containing *Cas9* nuclease gene, Puro$^{R}$ | |
| pLVX-*IRES-neo* | Lentiviral vector for bicistronic expression of a gene together with a neomycin-resistance marker | Clonetech, 632181 |
| pLentiCRISPRv2-gRNA-*DNAJC5* | pLentiCRISPRv2 with the gRNA targeting *DNAJC5* gene; Puro$^{r}$ | This study |
| pLVX-*DNAJC5-FLAG* | pLVX-*IRES-neo* vector which contains FLAG-tagged *DNAJC5* gene; Neo$^{R}$ | This study |
| pLVX-*DNAJC5*$^{S10A\text{-}S34A}$-*FLAG* | pLVX-*IRES-neo* vector which contains FLAG-tagged *DNAJC5* gene mutated on phosphorylation sites; Neo$^{R}$ | This study |
| pLVX-*DNAJC5*$^{H43Q}$-*FLAG* | pLVX-*IRES-neo* vector which contains FLAG-tagged *DNAJC5* gene mutated on Hsc70/Hsp70 binding site; Neo$^{R}$ | This study |
| pLVX-*DNAJC5*$^{L115R}$-*FLAG* | pLVX-*IRES-neo* vector which contains FLAG-tagged *DNAJC5* gene with L115R mutation in the cysteine-string domain; Neo$^{R}$ | This study |
| pLVX-*DNAJC5*$^{\Delta L116}$-*FLAG* | pLVX-*IRES-neo* vector which contains FLAG-tagged *DNAJC5* gene with the deletion of L116 in the cysteine-string domain; Neo$^{R}$ | This study |

| | | |
|---|---|---|
| pLVX-*DNAJC5*$^{\Delta Jdomain}$-*FLAG* | pLVX-*IRES-neo* vector which contains FLAG-tagged *DNAJC5* gene with the deletion of the J-domain; Neo$^{R}$ | This study |
| DNAJC5-GFP | Mammalian expression plasmid for DNAJC5 with N-terminal GFP tag | [8] |

**Supplementary Table 2: Oligonucleotides for PCR amplification**

| Names | Primers (5' → 3') |
|---|---|
| ExoU-Gm_Fw | GTCCGGCTCCGGAGTCAC |
| ExoU-Gm_Rev | GCTGCAGCATTTCGCGCG |
| ExoU-BamHI | CCGGATCCCAAGGCGCTTGATCAGTGG |
| ExoU-XbaI | GGTCTAGATGTGAACTCCTTATTCCGCCAAG |
| DNAJC5-Fw | GGAGTGCTGGGATGACAGG |
| DNAJC5-Rev | CAGTCCCTGGGATCTACGG |

**Supplementary Table 3: List of primers used to amplify and sequence the gRNA-containing cassettes of each sample after the ExoU screen**

| Names | Primers (5' → 3')* |
|---|---|
| Uninfected_Fw | AATGATACGGCGACCACCGAGATCTACACATAGAGGCACACTCTTTCCC TACACGACGCTCTTCCGATCTTTGTGGAAAGGACGAAACACCG |
| Uninfected_Rev | CAAGCAGAAGACGGCATACGAGATTCTCCGGAGTGACTGGAGTTCAGA CGTGTGCTCTTCCGATCTACTTGCTATTTCTAGCTCTAAAAC |
| ExoU-infected-1_Fw | AATGATACGGCGACCACCGAGATCTACACTATAGCCTACACTCTTTCCCT ACACGACGCTCTTCCGATCTTTGTGGAAAGGACGAAACACCG |
| ExoU-infected-1_Rev | CAAGCAGAAGACGGCATACGAGATTTCTGAATGTGACTGGAGTTCAGA CGTGTGCTCTTCCGATCTACTTGCTATTTCTAGCTCTAAAAC |
| ExoU-infected-2_Fw | AATGATACGGCGACCACCGAGATCTACACTATAGCCTACACTCTTTCCCT ACACGACGCTCTTCCGATCTTTGTGGAAAGGACGAAACACCG |
| ExoU-infected-2_Rev | CAAGCAGAAGACGGCATACGAGATACGAATTCGTGACTGGAGTTCAGA CGTGTGCTCTTCCGATCTACTTGCTATTTCTAGCTCTAAAAC |
| ExoU-infected-3_Fw | AATGATACGGCGACCACCGAGATCTACACTATAGCCTACACTCTTTCCCT ACACGACGCTCTTCCGATCTTTGTGGAAAGGACGAAACACCG |
| ExoU-infected-3_Rev | CAAGCAGAAGACGGCATACGAGATAGCTTCAGGTGACTGGAGTTCAGA CGTGTGCTCTTCCGATCTACTTGCTATTTCTAGCTCTAAAAC |

*Red sequences denote i5 or i7 index. Blue sequences denote annealing sequence on TKOv3 vector

**Supplementary Table 4: Oligonucleotides to knock-out the *DNAJC5* gene**

| Names | Primers (5' → 3')* |
|---|---|
| gRNA-DNAJC5_Fw | **CACCG**GAGGCCGCAGAAGACAAACA |
| gRNA-DNAJC5_Rev | **AAAC**TGTTTGTCTTCTGCGGCCTC**C** |

* Bold letters represent overhangs for cloning in *Bsm*BI-digested plentiCRISPRv2

**Supplementary Table 5: Synthetic *DNAJC5-FLAG* gene sequence and its derivatives**

| Names | Sequence (5' → 3')* |
|---|---|

| | |
|---|---|
| *DNAJC5-FLAG* | GAATTCATGGACTACAAAGACCATGACGGCGATTATAAAGATCATGACA<br>TCGATTACAAGGATGACGATGACAAGGCAGACCAGAGACAGCGCTCAC<br>TGTCTACCTCTGGGGAGTCATTGTACCACGTCCTTGGGTTGGACAAGAA<br>CGCAACCTCAGATGACATTAAAAAGTCCTATCGGAAGCTTGCCTTGAAAT<br>ATCACCCCGACAAGAACCCCGACAACCCGGAGGCCGCGGACAAGTTTAA<br>GGAGATCAACAACGCGCACGCCATCCTCACGGACGCCACAAAAAGGAA<br>CATCTACGACAAGTACGGCTCGCTGGGTCTCTACGTGGCCGAGCAGTTT<br>GGGGAAGAGAACGTGAACACCTACTTCGTGCTGTCCAGCTGGTGGGCC<br>AAGGC**T**CTGTT**C**GT**G**TT**T**TGCGGCCTCCTCACGTGCTGCTACTGCTGCTG<br>CTGTCTGTGCTGCTGCTTCAACTGCTGCTGCGGGAAGTGTAAGCCCAAG<br>GCGCCTGAAGGCGAGGAGACGGAGTTCTACGTGTCCCCCGAGGATCTG<br>GAGGCACAGCTGCAGTCTGACGAGAGGGAGGCCACAGACACGCCGATC<br>GTCATACAGCCGGCATCCGCCACCGAGACCACCCAGCTCACAGCCGACT<br>CCCACCCCAGCTACCACACTGACGGGTTCAACTAAGGATCC |
| *DNAJC5-FLAG*$^{S10A-S34A}$ | GAATTCATGGACTACAAAGACCATGACGGCGATTATAAAGATCATGACA<br>TCGATTACAAGGATGACGATGACAAGGCAGACCAGAGACAGCGCTCAC<br>TGGCTACCTCTGGGGAGTCATTGTACCACGTCCTTGGGTTGGACAAGAA<br>CGCAACCTCAGATGACATTAAAAAGGCCTATCGGAAGCTTGCCTTGAAA<br>TATCACCCCGACAAGAACCCCGACAACCCGGAGGCCGCGGACAAGTTTA<br>AGGAGATCAACAACGCGCACGCCATCCTCACGGACGCCACAAAAAGGA<br>ACATCTACGACAAGTACGGCTCGCTGGGTCTCTACGTGGCCGAGCAGTT<br>TGGGGAAGAGAACGTGAACACCTACTTCGTGCTGTCCAGCTGGTGGGC<br>CAAGGC**T**CTGTT**C**GT**G**TT**T**TGCGGCCTCCTCACGTGCTGCTACTGCTGCT<br>GCTGTCTGTGCTGCTGCTTCAACTGCTGCTGCGGGAAGTGTAAGCCCAA<br>GGCGCCTGAAGGCGAGGAGACGGAGTTCTACGTGTCCCCCGAGGATCT<br>GGAGGCACAGCTGCAGTCTGACGAGAGGGAGGCCACAGACACGCCGAT<br>CGTCATACAGCCGGCATCCGCCACCGAGACCACCCAGCTCACAGCCGAC<br>TCCCACCCCAGCTACCACACTGACGGGTTCAACTAAGGATCC |
| *DNAJC5-FLAG*$^{H43Q}$ | GAATTCATGGACTACAAAGACCATGACGGCGATTATAAAGATCATGACA<br>TCGATTACAAGGATGACGATGACAAGGCAGACCAGAGACAGCGCTCAC<br>TGTCTACCTCTGGGGAGTCATTGTACCACGTCCTTGGGTTGGACAAGAA<br>CGCAACCTCAGATGACATTAAAAAGTCCTATCGGAAGCTTGCCTTGAAAT<br>ATCAGCCCGACAAGAACCCCGACAACCCGGAGGCCGCGGACAAGTTTA<br>AGGAGATCAACAACGCGCACGCCATCCTCACGGACGCCACAAAAAGGA<br>ACATCTACGACAAGTACGGCTCGCTGGGTCTCTACGTGGCCGAGCAGTT<br>TGGGGAAGAGAACGTGAACACCTACTTCGTGCTGTCCAGCTGGTGGGC<br>CAAGGC**T**CTGTT**C**GT**G**TT**T**TGCGGCCTCCTCACGTGCTGCTACTGCTGCT<br>GCTGTCTGTGCTGCTGCTTCAACTGCTGCTGCGGGAAGTGTAAGCCCAA<br>GGCGCCTGAAGGCGAGGAGACGGAGTTCTACGTGTCCCCCGAGGATCT<br>GGAGGCACAGCTGCAGTCTGACGAGAGGGAGGCCACAGACACGCCGAT<br>CGTCATACAGCCGGCATCCGCCACCGAGACCACCCAGCTCACAGCCGAC<br>TCCCACCCCAGCTACCACACTGACGGGTTCAACTAAGGATCC |

* Bold letters represent the mutations to prevent the Cas9 endonuclease from cleaving the gene. Red letters represent the S10A, S34A and H43Q mutations. Green letters are *Eco*RI and *Bam*HI restriction sites. Yellow letters denote the 3X-*FLAG* sequence.

**Supplementary Table 6: Oligonucleotides for site-directed mutagenesis**

| Names | Primers (5' → 3')* |
|---|---|
| DNAJC5_L115R_F | CTGTTCGTGTTTTGCGGCCGCCTCACGTGCTGCTACTGC |
| DNAJC5_L115R_R | GCAGTAGCAGCACGTGAGGCGGCCGCAAAACACGAACAG |
| DNAJC5_ΔL116_F | CTGTTCGTGTTTTGCGGCCTCACGTGCTGCTACTGCTGCTGC |
| DNAJC5_ΔL116_R | GCAGCAGCAGTAGCAGCACGTGAGGCCGCAAAACACGAACAG |
| DNAJC5_ΔJdomain_F | CCAGAGACAGCGCTCACTGTCTACCGGTCTCTACGTGGCCGAGCAGTTTGG |
| DNAJC5_ΔJdomain_R | CCAAACTGCTCGGCCACGTAGAGACCGGTAGACAGTGAGCGCTGTCTCTGG |

* Red letters represent the mutated or inserted nucleotides
